# Tabula Sapiens reveals the non-coding RNA landscape across 22 human organs and tissues

**DOI:** 10.64898/2026.03.23.713770

**Authors:** Jaeyoon Lee, Madhav Mantri, Kavita Murthy, Luise A. Seeker, George Crowley, Robert C. Jones, Tabula Sapiens Consortium, Stephen R. Quake

**Author notes:** A list of authors and their affiliations appears in the supplementary materials.

## Abstract

The biological significance of non-coding RNAs has been increasingly appreciated as their roles in various cellular processes are uncovered. However, single-cell transcriptomic profiling of human samples has focused primarily on protein-coding genes by targeting polyadenylated RNA transcripts, leaving the expression patterns of non-coding RNA underexplored. Here, we expand Tabula Sapiens to the non-coding transcriptome with single-cell and single-nucleus total RNA sequencing across 22 human organs and tissues. By simultaneously profiling both polyadenylated and non-polyadenylated transcripts, the resulting dataset enables joint analysis of the protein-coding and non-coding transcriptomes at single-cell and subcellular resolution. Using these data, we assessed the cell type specificity of non-coding genes and found that a greater proportion of non-coding genes are differentially expressed by single cell types compared to protein-coding genes. We then compared single-cell and single-nucleus data from the same samples to infer subcellular localization patterns, revealing cell type-dependent nuclear and cytoplasmic enrichment of specific non-coding RNAs. Next, we showed that tRNA repertoires are cell type-specific and that this specificity is not simply explained by differences in codon usage across cell types. Finally, we characterized dynamic expression patterns of non-coding RNAs across the cell cycle and senescence-associated cell states, identifying non-coding genes with putative roles in cell division and growth arrest. Our work establishes a resource for investigating the landscape of non-coding RNAs across a diverse set of human tissues and cell types.

## Introduction

Single-cell transcriptomics has created powerful reference tools for examining human cell biology (*1–7*). However, previous large-scale molecular cell atlas efforts including prior Tabula Sapiens experiments (*1*, *2*) have mostly relied on poly(A) capture, which preferentially profiles protein-coding messenger RNAs (mRNAs) while systematically undersampling non-coding RNAs (ncRNAs). Since ncRNAs constitute the vast majority of the human transcriptome (*8*) and encompass diverse biotypes with broad cellular functions (*9*, *10*), this technical bias yields an incomplete view of the transcriptomic programs underlying cellular identity and physiology.

Here, we extend Tabula Sapiens to ncRNAs by applying the *in vitro* polyadenylation strategy of TotalX (*11*) to a broad range of human organs and tissues. Compared with standard poly(A) capture-based protocols, this approach retains protein-coding information while substantially increasing detection of ncRNAs, including long non-coding RNAs (lncRNAs), transfer RNAs (tRNAs), small nuclear RNAs (snRNAs), small nucleolar RNAs (snoRNAs), and microRNAs (miRNAs). We additionally profiled total RNA within single nuclei (*12*) to provide complementary, compartment-specific views of RNA biology. The resulting dataset enables joint analyses of the coding and non-coding transcriptomes with single-cell and subcellular resolution at whole-organism scale.

To demonstrate the utility of this dataset, we performed integrative analyses that leverage its ncRNA enrichment, single-cell resolution, and broad coverage of tissues and cell types. First, we identified cell type-specific ncRNAs across the atlas and compared the relative specificities of different biotypes, finding that ncRNAs are major determinants of cellular identity. Next, we compared the single-cell and single-nucleus portions of the dataset to quantify nuclear-cytoplasmic partitioning of ncRNA transcripts, which may provide insight into the functions of the many ncRNA genes that remain poorly understood. We also examined differential tRNA gene expression patterns across cell types and observed that tRNA repertoires are cell type-specific independent of codon usage, though factors besides variation in codon usage likely contribute to their regulation. Finally, by characterizing expression across cell-cycle stage and senescence, we identified non-coding genes associated with cell division and growth arrest.

## Results

### Joint profiling of protein-coding and non-coding RNA in human tissues

We applied single-cell and single-nucleus total RNA sequencing (*11*, *12*) to 22 distinct tissues (**Methods**) collected from a 50-year-old male human donor of European descent with known medical background (**Supplementary Information**), hereafter referred to as TSP33. The resulting dataset comprises 27 whole-cell and 21 nuclear samples split across the 22 tissues, 11 of which were profiled with both modalities (**Fig. 1A**). After combined *in vitro* and *in silico* depletion of highly abundant ncRNA species such as *RN7SK*, *RN7SL*, and the ribosomal RNAs, 105,471 cells and 121,398 nuclei passed quality control filtering (**Methods**). At the bulk level, ncRNAs comprised 18.7% of all detected transcripts across the entire dataset, representing a substantial enrichment across ncRNA biotypes compared to previous Tabula Sapiens data (**fig. S1A-B**).

**Figure 1.**
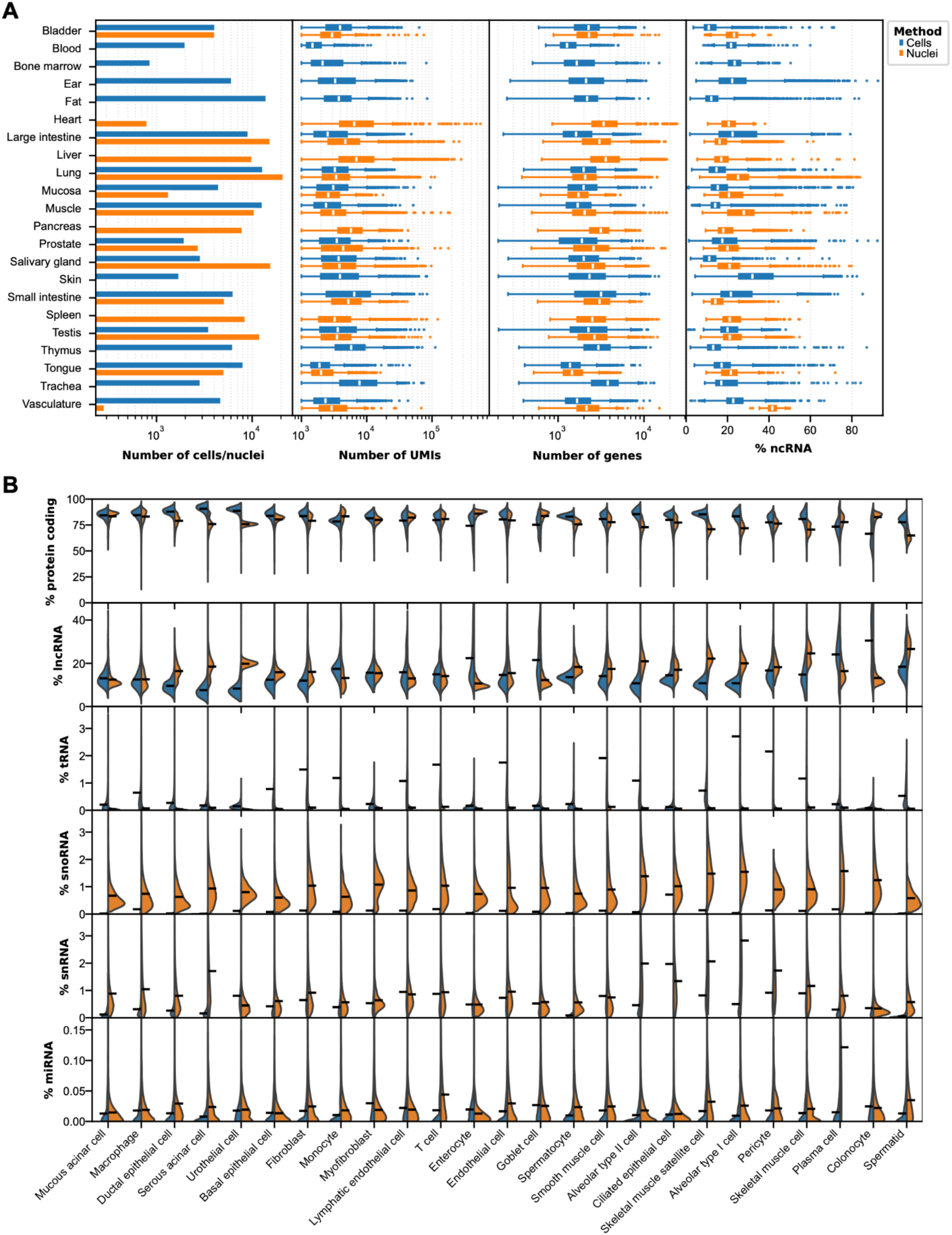
Dataset overview. A) Metadata and quality control metrics by sample. Bar chart showing the number of cells (blue) and nuclei (orange) profiled in each sample. Box plots showing the total number of UMIs detected, number of genes, and % of UMIs that mapped to ncRNA genes in each sample. B) Violin plots showing the distributions of transcript biotype proportions per cell type, split by modality. Only cell types represented by ≥100 cells and ≥100 nuclei in the dataset are included. Black line segments indicate the mean value for each half-violin.

Since the cell type specificities of protein-coding genes are well-characterized, we used the mRNA fraction of the TSP33 total RNA sequencing data to annotate 79 distinct cell types by applying large language model tools (*13*) and inspecting the expression of known cell type marker genes (**Methods**). We then examined the ncRNA fraction of the dataset and observed cell type-dependent differences in the proportion of transcripts in each biotype (**Fig. 1B**). In general, the single-cell data showed greater enrichment for tRNAs while the single-nucleus data were relatively enriched for snoRNAs and snRNAs (**fig. S1C**), consistent with the expected subcellular compartmentalization of these biotypes. The differences in biotype abundances between the two modalities were also cell type-dependent, with particularly stark differences for lncRNAs, which were more abundant in whole cells for some cell types and in nuclei for others. These quantifications of ncRNA abundances indicate that our dataset resolves expression patterns of ncRNAs at cell type resolution, with the two modalities capturing complementary aspects of RNA biology.

To assess concordance with previous Tabula Sapiens results, we integrated the TSP33 TotalX data with Tabula Sapiens 2.0 and compared matched cell populations profiled with the different protocols (**Methods**). The expression patterns of protein-coding genes were highly concordant between TSP33 and previous Tabula Sapiens donors (**fig. S2A-C**), demonstrating that TotalX recovered protein-coding information similarly to methods used previously in Tabula Sapiens 2.0. Non-coding genes, which include polyadenylated lncRNAs and other ncRNAs that may be incidentally captured by poly(A) capture-based methods through mispriming, were also highly concordant as a whole (**fig. S2D-E**), indicating that ncRNA expression patterns are broadly conserved across individuals. However, short nuclear-enriched ncRNA biotypes such as snoRNAs and snRNAs exhibited reduced concordance (**fig. S2F**), likely reflecting the lower sensitivity of previous methods for detecting these RNAs and underscoring the value of TotalX and single-nucleus approaches for more comprehensive profiling of the non-coding transcriptome.

### Cell type specificity of ncRNAs

Many ncRNA genes are known to exhibit cell type-specific expression. In particular, multiple studies have found that lncRNAs may display greater tissue and cell type specificity than protein-coding genes (*14–17*). Similarly, tRNAs and snoRNAs, which play central roles in protein synthesis and ribosome biogenesis in all cells, include members with striking tissue specificity (*18–22*), suggesting pronounced underlying cell type specificity. Despite these findings, characterization of ncRNA cell type specificity in humans has largely been limited to assays that target only a narrow subset of ncRNAs or by the use of bulk tissue profiling. To address these limitations, we leveraged the single-cell resolution of our total RNA sequencing dataset to investigate the cell type specificity of genes across the entire transcriptome and in a broad range of human cell types.

We first applied standard differential gene expression analysis (DGEA) methods to identify differentially expressed genes (DEGs) for each cell type across our dataset (**Methods**). Due to the variation in detection rates for each biotype between cells and nuclei (**Fig. 1B**), we considered the single-cell and single-nucleus portions of our dataset separately to reduce biases that may arise from differences in the cellular and nuclear proportions of each cell type in our dataset. Many DEGs were detected with both modalities, but they each preferentially detected particular biotypes (**fig. S3A**); we aggregated the results from the two portions of the dataset for each cell type to capture information from both.

We identified numerous ncRNA DEGs for each annotated cell type in our dataset (**Fig. 2A, table S1**). Most ncRNA DEGs were not previously noted to be cell type-specific, but our data also confirmed previously known expression patterns, such as *PCAT19* in endothelial cells (*23*) and the myomiRs *hsa-miR-133b* and *hsa-miR-206* in skeletal muscle cells (*24*). Taken together, these observations of known and unknown cell type-specific protein-coding and non-coding genes confirm the utility of our dataset in identifying novel cell type-specific ncRNAs.

**Figure 2.**
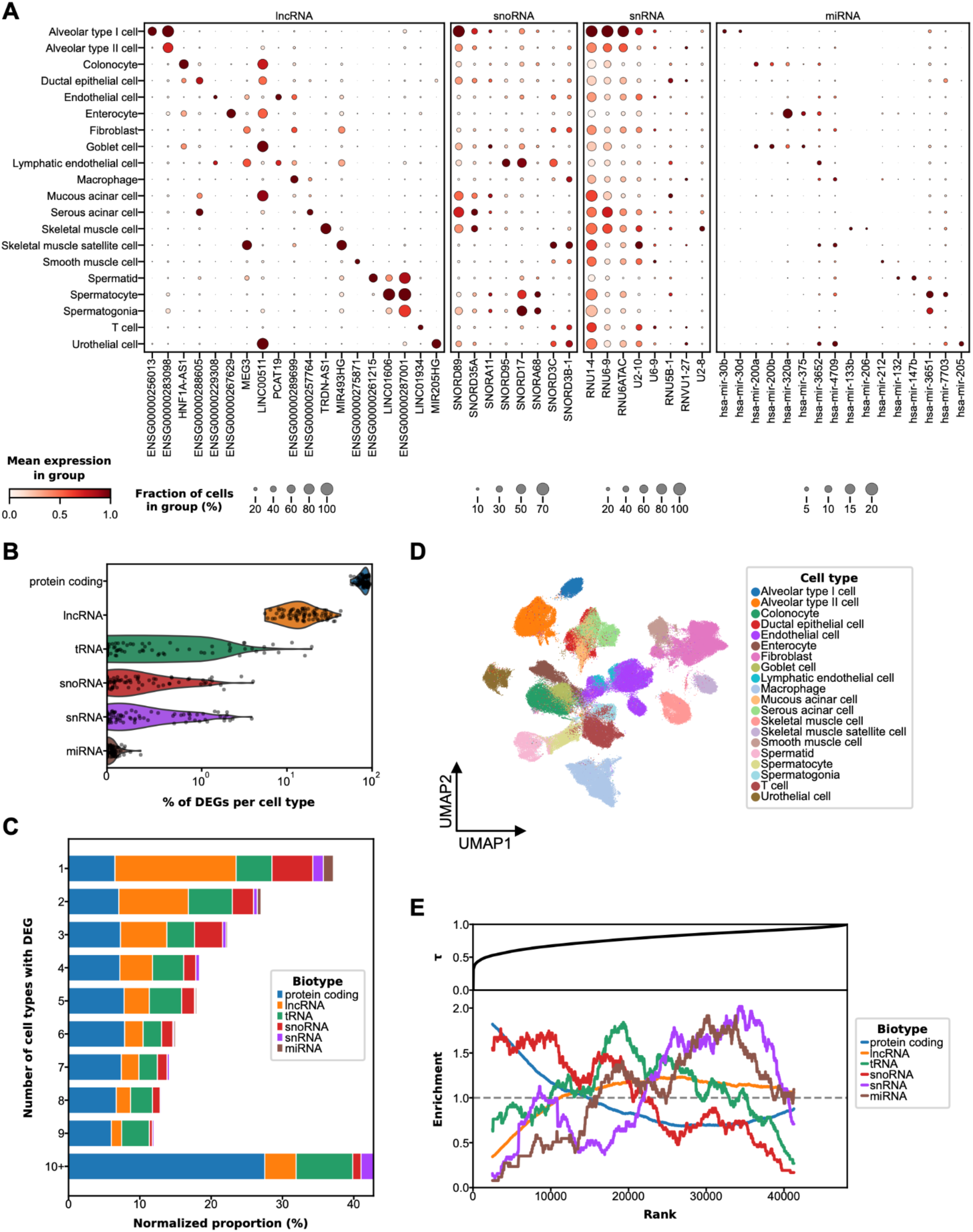
Cell type specificity of ncRNAs. A) Dot plots showing expression of representative cell-type specific lncRNA, snoRNA, snRNA, and miRNA genes. The top 20 cell types based on total cell and nuclei counts are shown. B) Violin plot showing the distribution of biotype proportions in the differentially expressed genes (DEGs) for each annotated cell type; points represent individual cell types. C) Stacked bar plot showing the normalized proportion of genes that were differentially expressed in a given number of cell types. Proportions are normalized by the number of annotated genes of the corresponding biotype. D) Uniform Manifold Approximation and Projection (UMAP) visualization of the top 20 cell types based on total cell and nuclei counts colored by cell type, clustered based on expression of ncRNA DEGs alone. E) Top: tissue specificity index *τ* plotted against gene rank, with genes ordered by increasing *τ*. Bottom: enrichment of each biotype within a moving window across the same gene ranking, normalized by the total number of genes of that biotype. An enrichment value of 1 indicates that the local proportion of genes from a given biotype equals its overall proportion across all annotated genes with ≥ 100 counts across the dataset.

To evaluate the relative cell type specificities of each biotype, we calculated the fraction of DEGs belonging to each biotype within each cell type (**Fig. 2B**). Overall, most DEGs were protein-coding genes, with this biotype making up 79.7% of DEGs for each cell type on average. However, protein-coding DEGs were frequently shared between multiple cell types, with only a small subset uniquely detected in single cell types. When normalized by the total number of annotated genes of each biotype in the human genome, a larger fraction of lncRNAs than protein-coding genes were DEGs uniquely in a single cell type (“unique DEGs”) (**Fig. 2C**). Similarly, the set of unique DEGs comprised greater proportions of ncRNA biotypes, with non-coding genes comprising 75.6% of these genes (**fig. S3B**). These results mirror previous studies suggesting that lncRNAs may be more cell type-specific than protein-coding genes (*17*, *25*). To test whether ncRNAs alone can define cell identity, we examined the expression patterns of ncRNA DEGs and found that these genes are sufficient to distinguish cell types (**Fig. 2D**).

For a more global view of the relative cell type specificities of all genes, we computed the specificity statistic *τ* for each gene (*26*) (**Methods**). This statistic increases with increasing specificity and has previously been used with Tabula Sapiens and other molecular cell atlases (*1*, *27*, *28*). Since the two modalities preferentially captured different biotypes (**Fig. 1B**), we calculated *τ* separately for cells and for nuclei and compared them for each gene (**fig. S4A**). The resulting values were moderately comparable (concordance correlation coefficient *p_c_* = 0.66), with some genes appearing more cell type-specific with one modality compared to the other. We surmised that these differences were due to increased dropout in the modality with less sensitivity to those genes, so we retained the lower value of *τ* to better reflect the expression profile of each gene.

We ranked genes by *τ* in ascending order and calculated the relative enrichments of major biotypes within a moving window across the ranks (**Fig. 2E, Methods**). Protein-coding genes were most enriched at the lowest *τ* values and decreased progressively with rank, in line with the vast number of proteins that are indispensable for all cells. snoRNAs showed a similar trend, concordant with their essential role in ribosome biogenesis. tRNAs were most enriched at intermediate ranks, consistent with both their requirement for protein synthesis and the extensive redundancy among tRNA gene copies. Other ncRNA biotypes were enriched mostly at higher ranks; lncRNAs, snRNAs, and miRNAs each contain more cell type-specific genes than protein-coding genes in proportion. The relative enrichment patterns of snRNAs and miRNAs were particularly distinct, with most members of these two biotypes exhibiting high cell type specificity. Previous works have suggested that some variant snRNA genes form alternative spliceosomes that may contribute to cell type-specific alternative splicing (*29–31*), and miRNAs are thought to be integral for cell type-specific post-transcriptional control of gene expression(*32–34*); our results uncover additional snRNA and miRNA genes that may be crucial determinants of cell type identity. These patterns were preserved when we quantified specificity using expression entropy instead of *τ* (**fig. S4B-C, Methods**), demonstrating robustness to the choice of specificity metric.

Besides identifying cell type-specific ncRNAs, we also found many ncRNAs that were widely expressed across many cell types. Ubiquitous lncRNAs are of particular interest because the roles of most lncRNAs remain elusive; such genes with unknown function may participate in essential cellular processes shared among all cells. Indeed, in the bottom 2% of lncRNAs ranked by *τ* (**table S2**), we find multiple genes shown to be indispensable across multiple human cell lines, such as *MALAT1*, *NEAT1*, and *GAS5* (*35*). Most of the other broadly expressed lncRNAs identified in our dataset currently have no known function but may also be necessary for cellular survival.

### Nuclear compartmentalization of ncRNAs

The subcellular localization of RNA has long been recognized as a critical determinant of RNA function. For mRNAs, spatial distribution within the cell is a crucial determinant of post-transcriptional gene regulation that varies across cell types and states (*36–39*), and mislocalization of specific transcripts has been implicated in a range of diseases (*40–42*). Similarly, multiple studies have shown that ncRNAs exhibit distinct subcellular localization patterns that vary with cell type and cellular state (*43–46*), underscoring the biological significance of ncRNA compartmentalization. However, studies of ncRNA localization in humans to date have been constrained by limited coverage of the non-coding transcriptome and reliance on bulk tissue measurements. Our dataset overcomes both of these constraints with matched single-cell and single-nucleus total RNA sequencing from the same samples, enabling transcriptome-wide estimates of nuclear compartmentalization in human cells at cell type resolution.

To investigate the nuclear compartmentalizations of transcripts, we first compared our nuclear and whole-cell data at a bulk level by calculating the log fold change in each gene’s proportional abundance between the two modalities after pseudobulking all samples of the same type. Although previous works have noted that such comparisons may not directly reflect differences in transcript counts between compartments and instead represent variation in RNA concentrations for each gene (*47*, *48*), we posit that these “relative enrichments” capture biologically relevant variation in RNA localization, processing, or stability between nuclei and whole cells. Moreover, since the kinetics of downstream RNA processes in each compartment likely depend on concentrations rather than absolute transcript counts, relative enrichments should provide a functionally meaningful view of subcellular transcript dynamics. Accordingly, variations in relative nuclear enrichment across genes or cell types likely correspond to biologically relevant differences in compartment-specific regulation and RNA activity.

We found that all analyzed biotypes comprised a mix of both nuclear-enriched and nuclear-depleted genes (**Fig. 3A, table S3**). Protein-coding and lncRNA genes were approximately evenly split between relative enrichment and depletion, consistent with the broad ranges of cellular processes in which they participate. In contrast, tRNA genes were largely depleted in nuclei while snRNA and snoRNA genes were mostly enriched in nuclei, consistent with the subcellular compartments where their canonical functions take place and the relative enrichments of these biotypes at the bulk level. However, we observed many exceptions in each of these biotypes. Since tRNAs and spliceosomal snRNAs undergo regulated nucleocytoplasmic transport during maturation (*49–51*), such exceptions may indicate variation in the kinetics of individual steps of their biogenesis pathways. Moreover, these ncRNAs may also perform noncanonical functions outside of their usual localizations, as suggested by previous works: tRNA fragments (tRFs) have been shown to act as epigenetic regulators in the nucleus (*52*), and snoRNAs can stabilize mRNAs in the cytoplasm (*53*). While miRNAs are known to comprise genes that are enriched in different compartments due to nuclear re-import (*54*), we found that a substantial majority were relatively enriched in the nucleus, which may result from the detection of pri– and pre-miRNAs that are yet to be exported from the nucleus in addition to mature miRNAs that may have been re-imported.

**Figure 3.**
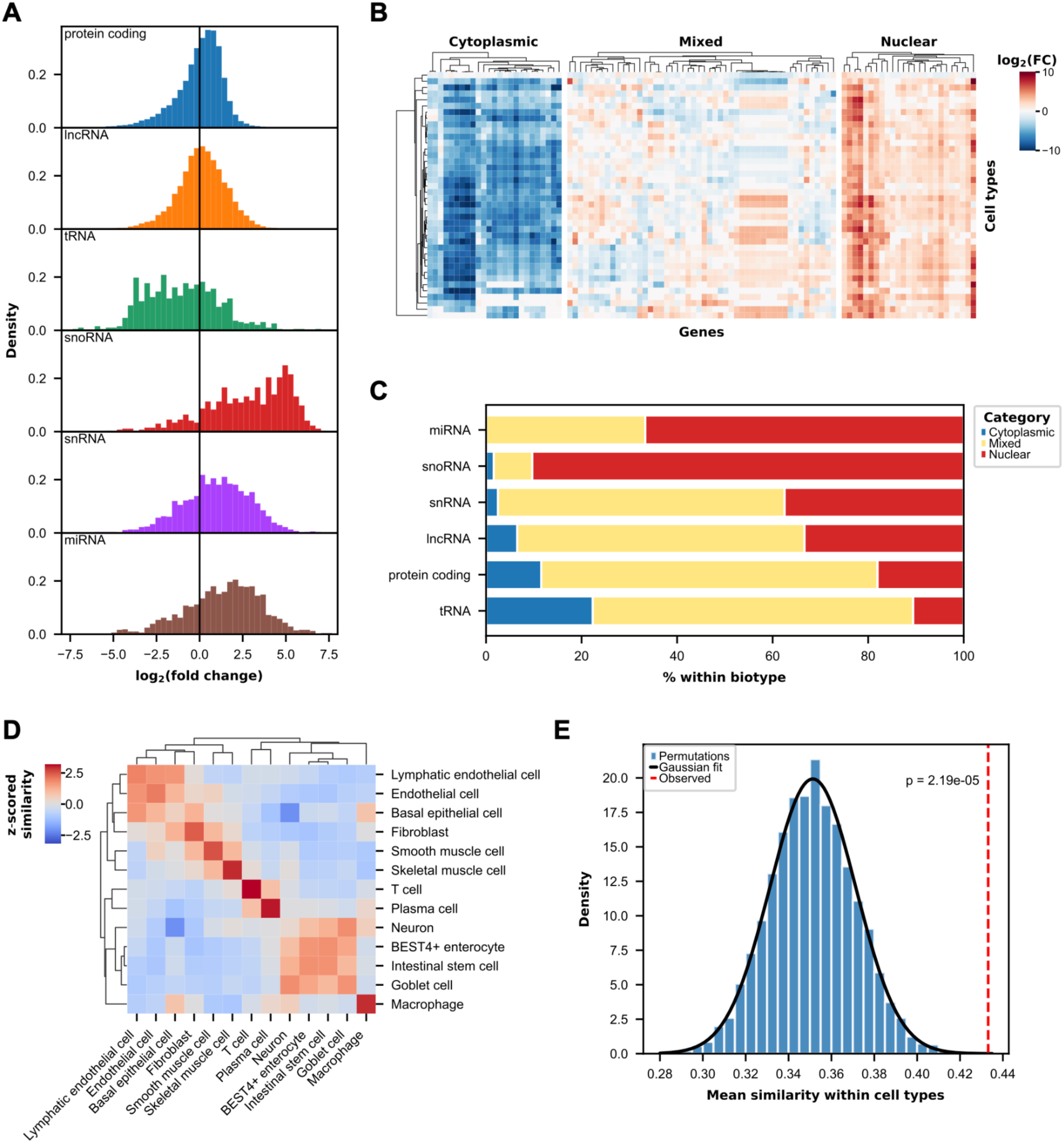
RNA nuclear compartmentalization across biotypes and cell types. A) Histograms showing the distributions of nuclear enrichment across genes in selected biotypes at the bulk level. B) Heat maps showing nuclear enrichment values for representative genes (columns) that are predominantly depleted in nuclei (left), display mixed enrichment across cell types (middle), or are predominantly enriched in nuclei (right). Rows correspond to cell types that are well-represented in both modalities. Rows and columns are clustered by Euclidean distance, as indicated by the dendrograms above and to the left. C) Stacked bar chart showing the proportions of each biotype that are predominantly cytoplasmic, display mixed enrichment across cell types, or are predominantly nuclear. D) Heat map showing mean z-scored cosine similarities of nuclear compartmentalization status for each gene across tissue-cell type combinations, aggregated at the cell type level. Only tissue-cell types represented by ≥50 cells and ≥50 nuclei were considered, and only cell types which passed these thresholds in at least two tissues were included. For self-correlations within a cell type, only pairs of distinct tissues were compared. Rows and columns are clustered by Euclidean distance, as indicated by the dendrograms above and to the left. E) Histogram showing the mean of per-cell type mean cosine similarities of nuclear compartmentalization status across 1,000 permutations of tissue-cell type labels, fitted to a normal distribution (black curve). The observed mean (red dashed line) is significantly higher than the permuted distribution (p = 2.19 ✕ 10^-5^).

We next investigated patterns of relative nuclear enrichments across different cell types. For the 41 cell types that were well-represented in our dataset with both single-cell and single-nucleus sequencing, we performed DGEA between modalities to identify genes that are significantly enriched or depleted in nuclei for each cell type (**Methods**). We then examined the nuclear enrichment values for each gene that showed significant differences between the modalities in at least 50% of the cell types (**Fig. 3B**, **table S4, Methods**). Of the 5,519 genes that passed these thresholds, 1,179 genes were “predominantly nuclear” (significant nuclear enrichment in >50% of cell types; none depleted) and 611 genes were “predominantly cytoplasmic” (significant nuclear depletion in >50% of cell types; none enriched). All biotypes besides miRNAs contained members in both sets (**Fig. 3C**), including lncRNAs with unknown function; the subcellular localizations of these genes may offer important insights into their diverse cellular roles and regulatory contexts (*43*). tRNAs, snoRNAs, and snRNAs were preferentially associated with the set that aligned with the location of their canonical functions, but we observed a small number of genes of each biotype in the opposite sets. For example, multiple Asp-GTC tRNA isodecoders were predominantly nuclear; these tRNAs may be exported less efficiently from the nucleus or re-imported more frequently to perform nuclear functions.

Most genes exhibited mixed compartmentalization patterns across cell types, showing nuclear enrichment in some and depletion in others (**Fig. 3C**). To assess cell type-specificity in these nuclear enrichment patterns, we calculated cosine similarities of compartmentalization status for each gene (enriched, depleted, or not significantly different in nuclei relative to whole cells) across tissue-cell type combinations that were well-represented in at least 2 distinct tissues with both modalities in our dataset (**Methods**). We then compared the mean of cosine similarities within cell types and between cell types (**Fig. 3D**). We observed greater similarities within the same cell type than across different cell types, although we also saw some concordance between closely related cell types, such as endothelial and lymphatic endothelial cells, smooth and skeletal muscle cells, and cell types of the intestinal epithelium. Permutation testing based on shuffled tissue-cell type labels confirmed that within-cell type similarities were significantly higher than expected by chance (**Fig. 3E, Methods**). Collectively, these results indicate that cell type-specific regulatory programs underlie the subcellular compartmentalization of diverse RNA classes.

### Cell type specificity of tRNA repertoires

tRNAs constitute the most numerous RNA biotype in the cell (*55*), concordant with their central role in protein synthesis. Although all cells require broad anticodon coverage in their tRNA repertoires to sustain translation, the relative abundances of specific tRNA genes vary between tissues and cell types (*18*, *19*, *56*, *57*). This variation may reflect differences in codon usage between cell types since translational efficiency depends on a precise balance between the tRNA anticodon abundance and the codon usage profiles of a cell: decoding is the rate limiting step of translation elongation (*58*), and the decoding rates of specific codons are correlated with the abundances of their corresponding anticodons (*59*, *60*). Indeed, previous works have shown that global translation rates in *E. coli* and yeast depend on the balance of anticodon supplies and codon usage (*61–63*). However, studies mapping the cell type specificity of tRNA repertoires and their alignment to codon usage profiles have been hindered by the inability to measure tRNA abundances with single-cell resolution or to simultaneously profile the codon usage in the same cells. Our dataset directly quantifies tRNA abundances in single cells while simultaneously capturing codon usage information based on mRNA abundances, enabling a unified analysis of tRNA supply and demand in the same cells.

We first searched for cell type-specific patterns in tRNA repertoires by examining Pearson correlations between mean tRNA abundance profiles of tissue-cell types with adequate tRNA coverage (**fig. S5A-B**, **Methods**). To mitigate the effect of mapping ambiguity between highly similar tRNA gene copies, we aggregated reads within each isodecoder family. We observed greater mean correlations within cell types in different tissues compared to across unrelated cell types (**Fig. 4A**). Permutation testing based on shuffling tissue-cell type labels showed that these differences were statistically significant for most analyzed cell types (**Fig. 4B, Methods**), indicating that tRNA profiles are broadly cell type-specific. Previous studies reporting cell type-specificity of tRNA repertoires have focused largely on neurons within mouse and human developmental brains (*56*); our results extend these findings to other cell types in adult human tissues through direct quantifications of tRNA abundances. We identified the tRNA gene families driving these patterns by performing DGEA across the cell types exhibiting significant specificity in their tRNA repertoires (**Fig. 4C**, **table S5, Methods**). In contrast to previous findings in brain neurons (*56*, *64*), we did not observe upregulation of the tRNA^Ala^(AGC) family in peripheral neurons, suggesting that neuronal tRNA regulation may be uniquely specialized in the central nervous system.

**Figure 4.**
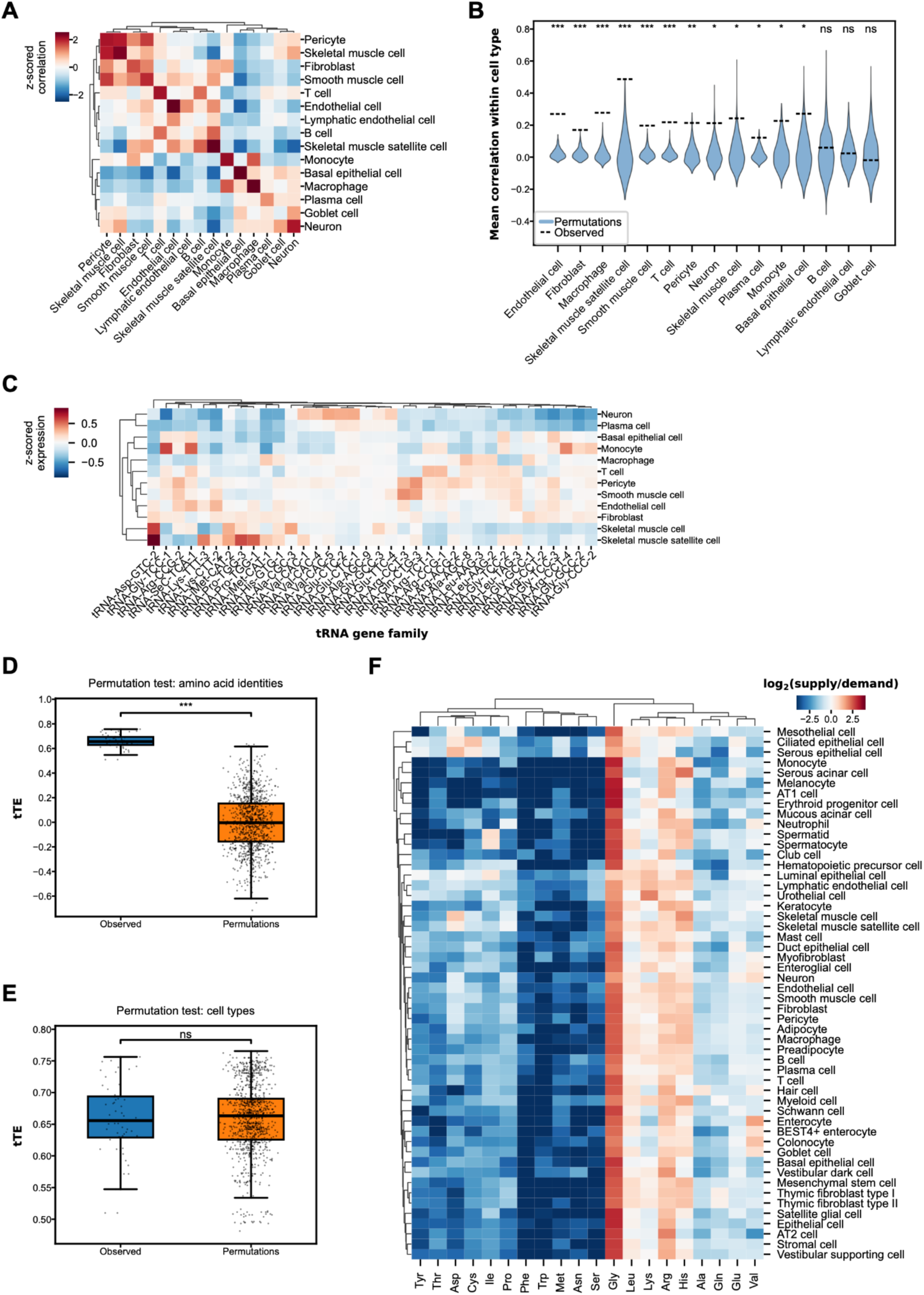
Cell type specificity of tRNA repertoires and balance with amino acid usage. A) Heat map showing z-scored mean Pearson correlations of tRNA repertoires across tissue-cell type combinations, aggregated at the cell type level. Only cell types that appeared in at least three tissues were included. For self-correlations within a cell type, only pairs of distinct tissues were compared. Rows and columns are clustered by Euclidean distance, as indicated by the dendrograms above and to the left. B) Violin plot showing the distributions of mean Pearson correlations within cell types grouped by tissue from 1,000 permutations of tissue-cell type labels. Dashed lines indicate the observed values, and cell types are arranged in order of decreasing significance. Asterisks display level of significance for each cell type: (ns) not significant, (*) *p* < 0.05, (**) *p* < 0.01, (***) *p* < 0.001. C) Heat map showing z-scored mean expression of the top five differentially expressed tRNA isodecoder subfamilies (columns) for cell type with significant specificity in their tRNA repertoires (rows). Rows and columns are clustered by Euclidean distance, as indicated by the dendrograms above and to the left. D) Box plots showing the distributions of observed and permuted theoretical translation efficiencies (tTEs) across cell types, with permutations generated by shuffling tRNA isotype identities. tTE was defined as the Spearman’s correlation between tRNA isotype supplies and amino acid demands inferred from protein-coding transcript abundances. Asterisks display level of significance: (***) *p* < 0.001. E) Box plots showing the distributions of observed and permuted tTEs across cell types, with permutations generated by shuffling tissue-cell type labels. The observed and permuted distributions were not significantly different (ns, not significant). F) Heat map showing the ratio of tRNA supply to demand across cell types. Rows and columns are clustered by Euclidean distance, as indicated by the dendrograms above and to the left.

We next investigated whether the observed cell type specificity of tRNA repertoires matched the cell type specificity of translational demands. For each cell type, we inferred the mean usage of each amino acid from the mRNA portion of our dataset (**Methods**). We then calculated the theoretical translation efficiency (tTE), defined as the Spearman’s correlation coefficient between the tRNA isotype abundance and amino acid usage profile, for each cell type (**Methods**). This metric has been used previously by combining separate datasets to show that human neurons exhibit greater translation efficiency compared to non-neuronal cell types, which were mostly comparable to one another (*64*). In agreement with these previous results, tTEs calculated from our dataset indicated well-matched tRNA supply and demand for most cell types, with no cell type exhibiting exceptionally high or low values (**fig. S5C**). Permutation testing based on shuffling amino acid identities in tRNA abundances revealed that the observed tTEs were significant for most cell types (**Fig. 4D, Methods**). However, observed tTEs were not elevated relative to values calculated from random pairings of tRNA repertoires and amino acid usages across different cell types (**Fig. 4E**, **Methods**). Taken together, these analyses suggest that while tRNA pools are globally well-aligned to translational demands, other factors besides translational demands shape tRNA repertoires in a cell type-specific manner.

Since tRNA isotype abundances are not finely adjusted to translational demand, different cell types likely exhibit varying degrees of misalignment between tRNA supply and amino acid usage. To assess this cell type dependence, we quantified supply-demand mismatches for each amino acid across cell types (**Fig. 4F**). Across most cell types, glycine and arginine tRNA isotypes were consistently over-supplied, whereas several others were consistently under-supplied, with tryptophan, phenylalanine, and asparagine tRNA isotypes showing the strongest depletion. Certain isotypes also showed coordinated patterns among closely related cell types. For example, isoleucine tRNAs were over-supplied in spermatocytes and spermatids, while valine tRNAs were over-supplied and lysine tRNAs under-supplied in the intestinal epithelium. These results point to potential cell type-specific vulnerabilities in tRNA supply that may constrain translational efficiency, as recently demonstrated across neuron subtypes in mice (*56*).

### ncRNAs dynamics throughout the cell cycle and in cells expressing senescence-associated transcripts

Single-cell mRNA sequencing has charted dynamic changes in mRNA expression patterns across a wide variety of cell states. In particular, molecular cell atlases have linked multiple protein-coding genes to cell-cycle phases (*65–67*) and cellular senescence programs (*68–70*). Specific ncRNAs have also been demonstrated to influence cell division and growth arrest through various mechanisms, such as altering chromatin structure, modulating mRNA stability and splicing, and acting as scaffolds or decoys for other functional molecules (*71–75*). However, many ncRNAs with roles in these processes may not yet have been identified due to the limited detection of ncRNAs with conventional single-cell transcriptomics methods. To address this gap, we examined the TSP33 total RNA sequencing dataset to identify ncRNAs with putative roles in cell division and cellular senescence.

For cell types that are known to divide, we first classified each cell and nucleus in our dataset as being in G1, S, or G2/M phase by using an established list of cell-cycle markers (*66*) (**Methods**). We then identified phase-specific shifts in RNA biotypes by calculating the ratio of their average proportion in each cell-cycle phase compared to overall average for each cell type (**Fig. 5A**). The relative abundances of most RNA biotypes remained stable across cell-cycle stages, with slight reductions in tRNAs, snRNAs, and snoRNAs from G1 to S phase, followed by slight increases toward G2/M in most cell types. These fluctuations may reflect subtle shifts in the activity of these biotypes, which are primarily involved in protein synthesis and ribosome biogenesis, while the cell is focused on DNA synthesis. We also observed a slight decrease in lncRNA abundance accompanied by a corresponding increase in miRNA abundance between the S and G2/M phases.

**Figure 5.**
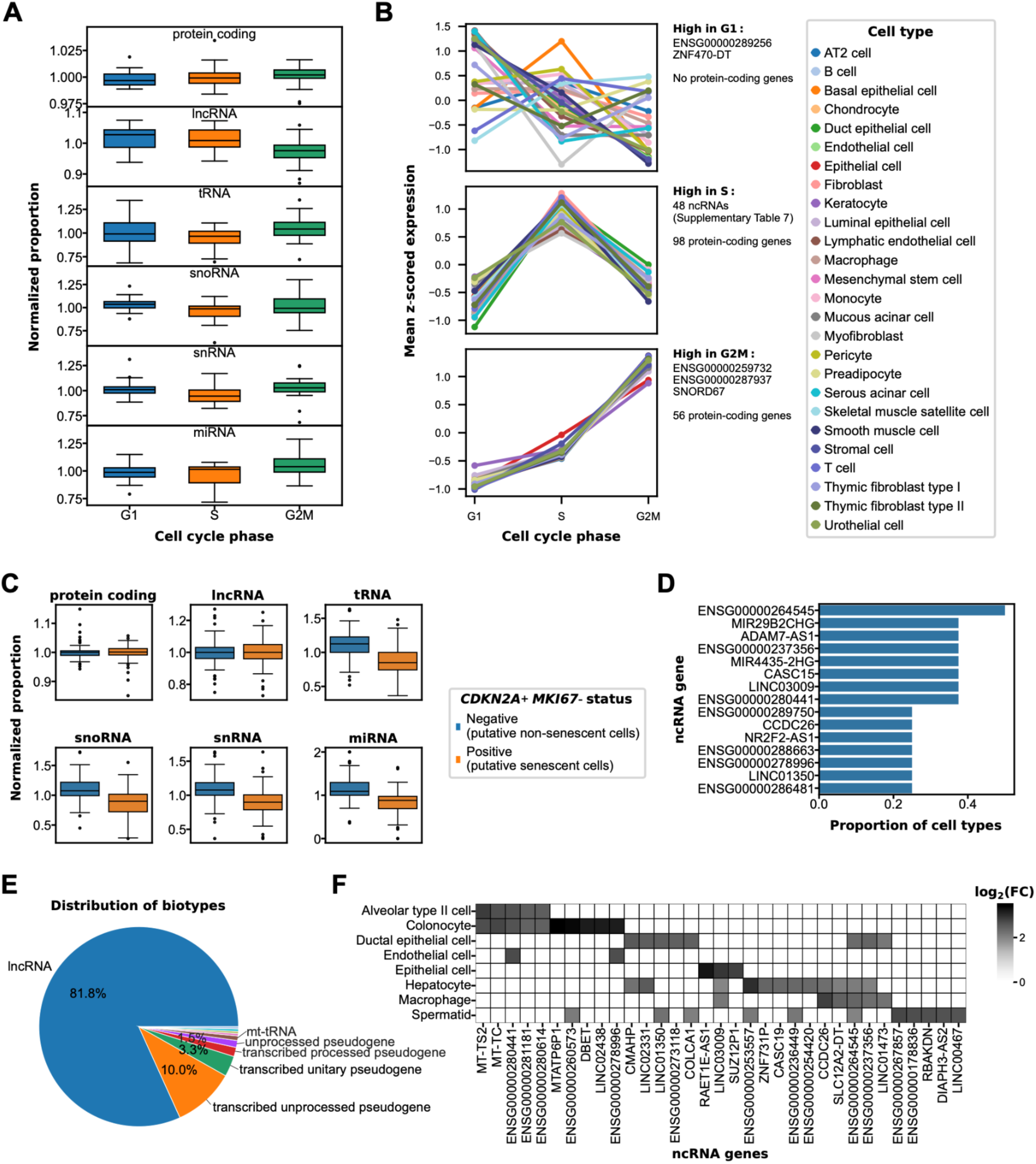
ncRNAs dynamics throughout the cell cycle and in cells expressing senescence-associated transcripts. A) Box plots showing the mean normalized proportions of RNA biotypes across cell types in each cell-cycle phase. Each plot was normalized to the mean of the per-cell type abundance across all phases. B) Plots showing mean z-scored expression of genes consistently upregulated in G1 (top), S (middle), and G2/M (bottom) phases within each cell type. The number of protein-coding genes and the identities of non-coding genes in each group are shown to the right of each plot. C) Box plots showing the mean normalized proportions of RNA biotypes across cell types for putative non-senescent and putative senescent cells. Each plot was normalized to the mean of the per-cell type abundance across both states. D) Bar plot of the top 15 senescence-associated ncRNA genes, ranked by the proportion of cell types in which each gene is enriched in *CDKN2A*+ *MKI67*-cells relative to all other cells. E) Pie chart showing the distribution of RNA biotypes enriched in *CDKN2A*+ *MKI67*-cells as compared to others. Long non-coding RNAs constitute the vast majority of detected species (81.8%), followed by smaller fractions pseudogene-derived transcripts and other annotated biotypes. F) Heat map of log fold-change values for top five non-coding RNA enriched in *CDKN2A*+ *MKI67*-cells across broad cell types. Rows represent annotated broad cell populations, columns represent ncRNA genes, and grayscale intensity reflects the magnitude of differential expression, revealing cell type-specific enrichment patterns.

We next sought to identify specific ncRNA genes with potential roles in cell division by examining phase-dependent expression patterns within each cell type. For the whole cell and nuclear portions of our dataset separately, we calculated mean expression profiles of gene-cell type pairs across the three cell-cycle phases and clustered these profiles into six groups with similar phase-specific expression dynamics (**fig. S6A**, **table S6, Methods**). Most genes exhibited cell type-specific phase patterns and therefore fell into different groups across cell types. To identify candidate genes that may be universally involved in cell division, we searched for those consistently assigned to the same cluster across cell types in which they were detected (**table S7**, **Methods**). This approach recovered all markers used to annotate cell-cycle phases in both whole cells and nuclei, indicating that our method detects upregulated genes in the correct phases (**fig. S6B**). In addition, we identified multiple ncRNA genes that were consistently upregulated in each phase (**Fig. 5B**). In particular, some lncRNA genes were among the most consistently upregulated genes in the S phase, such as *ENSG00000244332* and *TALAM1* in whole cells (**fig. S6C**) and *ENSG00000244332*, *ENSG00000256139*, and *ENSG00000289475* in nuclei (**fig. S6D**). One of these genes, *ENSG00000244332,* sense-overlaps the protein-coding gene *HELLS*, so our detection of this gene may arise from misassignment of *HELLS* transcripts. Nevertheless, *HELLS* is well-known to peak in S phase (*76*, *77*) and is a commonly used S phase marker (*66*) (including in our analysis), so the apparent upregulation of *ENSG00000244332* during S phase further validates our approach for identifying other replication-associated transcripts.

We also used the protein-coding portion of the TSP33 total RNA sequencing dataset to identify cells expressing senescence-associated transcripts in our dataset (**Methods**). To uncover senescence-associated changes in ncRNA expression patterns, we compared biotype abundances between *CDKN2A+ MKI67*-cells and other cells across cell types (**Fig. 5C**, **Methods**). We found that, for most cell types, *CDKN2A+ MKI67*-cells have slightly reduced snoRNA, snRNA, and miRNA content as compared to other cells of the same cell types, concordant with previously observed disruptions in ribosome biogenesis and post-transcriptional gene regulation in cellular senescence (*78–81*).

We then searched for senescence-associated changes in specific ncRNA genes across different cell types by performing DGEA between *CDKN2A+ MKI67*– and other cells within tissue-cell type groups (**Methods**). We identified a total of 269 putative senescence-associated ncRNA genes enriched in *CDKN2A+ MKI67*-cells across cell types (**table S8**). As was the case for protein-coding genes (*1*), no ncRNA gene was universally enriched in *CDKN2A+ MKI67*-cells for all analyzed cell types (**Fig. 5D**). Among the ncRNAs enriched in multiple cell types, *MIR4435-2HG* and *MIR29B2CHG* were associated with senescence for two cell types. Although most of the senescence-associated ncRNAs identified by our analysis have not been previously characterized in cellular senescence, the miRNA that is hosted in the *MIR29B2CHG* gene was recently shown to increase with aging in bone tissues and in senescent mesenchymal stroma (*82*), suggesting that our analysis might shed light on other non-coding features of senescence. Biotype analysis of senescence-associated ncRNA genes revealed lncRNA to be the most abundant class (**Fig. 5E**), extending previous studies that identified various lncRNA genes as crucial regulators of cellular senescence (*75*, *83*, *84*). Enrichment patterns of specific ncRNA genes varied by cell type (**Fig. 5F**), suggesting that diverse ncRNA genes participate in heterogeneous, cell type-dependent senescence-associated programs.

## Discussion

In this work, we present a comprehensive single-cell and single-nucleus total RNA sequencing atlas across 22 distinct human organs and tissues, broadening the scope of the Tabula Sapiens dataset to include the non-coding transcriptome. By capturing both the coding and non-coding transcriptomes across diverse cell types and states, the resulting dataset enables integrative analyses of previously inaccessible aspects of RNA biology. The vignettes included in this study highlight the value of these data in probing underexplored topics that are currently of active interest. As a whole, they illuminate the regulatory landscapes of ncRNA expression and distinguish functionally promising ncRNA candidates from a background of numerous poorly understood ncRNA genes, suggesting potentially fruitful directions for future studies with orthogonal approaches.

Our survey of ncRNA expression profiles across cell types identified both cell type-specific and ubiquitous ncRNA genes (**Fig. 2**). Although most ncRNA genes remain poorly characterized, these patterns identify ncRNA genes that may be functionally important and merit further study. Breadth of expression may offer clues to the functions of these genes: cell type-specific ncRNAs may contribute to processes specific to their host cell types while ubiquitous ncRNAs may participate in core functions broadly essential for cellular survival. In addition, our dataset reveals differing cell type specificity across RNA biotypes. Notably, several ncRNA biotypes exhibited greater cell type specificity than that of protein-coding genes, emphasizing the importance of defining how ncRNAs shape cellular identity.

The matched single-cell and single-nucleus portions of our dataset enabled a broad examination of subcellular localization of ncRNA transcripts at cell type resolution (**Fig. 3**). Our analysis showed that nuclear compartmentalization varies widely between genes and in a cell type-dependent manner, underscoring the biological significance of subcellular localization, including for ncRNAs. In addition, we observed that many ncRNA genes were consistently enriched or consistently depleted in nuclei across the profiled cell types. Interestingly, we found examples of ncRNA transcripts that were enriched in compartments opposite to those expected from their known functions; future work could elucidate whether these genes have noncanonical biological roles or if their localization is affected by inefficiencies in their biogenesis pathways.

Our examination of tRNA abundance profiles revealed that human tRNA repertoires are broadly cell type-specific (**Fig. 4**). While previous studies reported striking differences in the tRNA repertoires of neurons compared to those of non-neuronal cells, we show for the first time that this specificity extends to other cell types through direct quantification of tRNA expression. Furthermore, we show that while tRNA repertoires are globally well-aligned to translational demands, they are not finely tuned to minor cell type-dependent variations in amino acid demands. Instead, other regulatory mechanisms seem to more strongly drive tRNA expression patterns in a cell type-specific manner, and future functional experiments could elucidate the pathways involved.

By analyzing the dynamics of ncRNA expression as a function of cell state, we identified candidate genes with putative roles in cell division and growth arrest (**Fig. 5**). In particular, several poorly-characterized lncRNA genes were consistently upregulated in S phase across many cell types, indicating that these genes may participate in DNA replication. Furthermore, we found heterogeneous sets of ncRNA genes that were upregulated in cells expressing senescence-associated transcripts across diverse cell types, suggesting that various ncRNAs are involved in a wide range of senescence-associated phenotypes. Understanding the diversity of non-coding features of cells expressing senescence-associated molecules expands the landscape of actionable targets beyond protein-coding genes alone, offering promising avenues for developing targeted senolytics tailored to age-related diseases in specific tissues and cell types. More broadly, future targeted studies may clarify the mechanisms by which non-coding genes influence the overall regulation of cell cycling and cellular senescence, and extending similar analyses to other aspects of cell state could identify ncRNA genes involved in other processes of interest.

A key limitation of this study is the single-donor nature of the dataset. Although the TotalX data from this donor were broadly concordant with previous Tabula Sapiens data (**fig. S2**), profiling additional donors would increase power and confidence for detecting gene-level effects. Furthermore, we expect that incorporating donors with diverse genetic backgrounds and health profiles will enable deeper insights into ncRNA biology, paralleling findings from mRNA-focused atlases on age-, sex-, and disease-related variation in the protein-coding transcriptome. Beyond expansion of the dataset, improvements in methodology could enable deeper profiling of RNAs that were missed in this study, and enhanced computational methods could improve quantification of short and highly redundant ncRNAs, such as miRNAs and tRNAs.

Our work establishes a landmark resource for surveying the human non-coding RNA landscape. The analyses presented in this study demonstrate its applicability to a diverse array of topics and suggest multiple follow-up functional studies. As demonstrated by the impact of previous Tabula Sapiens datasets, we anticipate that this resource will be broadly useful for additional in-depth exploratory analyses in many areas of RNA biology.

## Acknowledgements

We thank the anonymous donor and his family for giving both the gift of life and the gift of knowledge through their generous donations. We thank Donor Network West for procuring the organs and tissues from the donor for this project. We are also grateful to the UCSF Liver Center (funded by NIH P30DK026743) for assistance with the liver samples. We thank members of the Quake lab for helpful discussion and comments. We also thank Dr. Alina Isakova for assistance with TotalX protocols and Dr. Mauricio Aguilar Rangel for technical advice regarding the tRNA analysis in this study.

## Funding

This project has been made possible in part by grant nos. 2019-203354, 2020-224249, 2021-237288, 2021-006486, 2022-316725. from the Chan Zuckerberg Initiative DAF; an advised fund of Silicon Valley Community Foundation; and by support from the Chan Zuckerberg Biohub San Francisco.

## Author contributions

J.L., S.R.Q., and the Tabula Sapiens Consortium conceptualized and designed experiments. The Tabula Sapiens Consortium collected live-cell and fresh-frozen samples and prepared dissociated single-cell suspensions. J.L., K.M., and G.C. prepared single-cell total RNA sequencing libraries with assistance from the Tabula Sapiens Consortium. L.A.S. and K.M. prepared nuclear suspensions from fresh-frozen samples. J.L. and K.M. prepared single-nucleus total RNA sequencing libraries with support from M.M on protocols. J.L and M.M. analyzed data with contributions from G.C on pre-processing and R.C.J. on cell type specificity analysis. J.L., M.M., and L.A.S. wrote the manuscript, and all authors contributed to revisions. S.R.Q. supervised the project.

## Competing interests

The authors declare no competing interests.

## Data, code, and materials availability

● The code used for the analysis and figure generation is available as a GitHub repository (https://github.com/jaeyoonl/Tabula-Sapiens-total-RNA/).
● Gene counts and metadata are publicly available from figshare (https://doi.org/10.6084/m9.figshare.31827436).
● The raw data files will be available from a public AWS S3 bucket upon publication.
● To preserve the donor’s genetic privacy, we require a data transfer agreement to receive the raw sequence reads.
● Any additional information required to reanalyze the data reported in this paper is available from the lead contact upon request.

## Materials and Methods

### Organ and tissue procurement

Organ and tissue procurement procedures followed previous methods from the Tabula Sapiens project (*1*, *2*). In brief, organs and tissues were procured through collaboration with Donor Network West (DNW, SanRamon, CA, USA) after obtaining records of first-person authorization and consent from the family members of the donor. Protocols were approved by DNW’s internal ethics committee (project STAN-19-104), the medical advisory board, and the Institutional Review Board at Stanford University which determined that this project does not meet the definition of human subject research as defined in federal regulations 45 CFR 46.102 or 21 CFR 50.3.

Each tissue was collected and transported on ice to tissue expert laboratories at Stanford and UCSF using a private courier service to limit the time between procurement and initial tissue preparation to less than one hour.

### Preparation of tissue samples and cell suspensions

Protocols for tissue preservation, tissue dissociation, and staining of cell suspensions for all tissues are described in the Tabula Sapiens publications (*1*, *2*) with the exception of the buccal mucosa sample, which was processed in the same way as the skin sample.

### Single-cell total RNA sequencing with TotalX

Protocols for single-cell total RNA sequencing with TotalX are described in the TotalX publication (*11*).

### Nuclei extraction and fixation

Fresh frozen human tissue was processed using the Sigma-Aldrich Nuclei Isolation Kit (NUC201-1KT). Tissue aliquots (mean 140 mg; range 19.3–689.5 mg) were finely chopped in a cryostat at −20 °C, transferred to RNase-free 1.5 ml LoBind tubes (Eppendorf, 0030108051), and stored in liquid nitrogen until use. All steps were performed on ice unless stated otherwise.

Tissue was lysed in 200 µL lysis buffer (PURE buffer; Sigma-Aldrich NUC 201 kit) supplemented with 0.1 M DTT (Thermo Fisher Scientific, A39255; 2 µL per sample), 10% Triton X-100 (2 µL per sample), and RNase inhibitor (Roche Protector RNase Inhibitor, 40 U/µL; Millipore Sigma, 3335402001; 220 U per sample), and homogenized using RNase-free disposable pestles (Fisherbrand, 12-141-364).

Lysates were mixed with 360 µL sucrose cushion and filtered through a 30 µm cell strainer (Sysmex Partec sterile and single-packed CellTrics, 04-004-2326). The sucrose cushion (560 µL per sample) was prepared fresh from PURE 2 M sucrose solution and sucrose cushion buffer (Sigma-Aldrich, NUC 201), supplemented with 0.1 M DTT and RNase inhibitor. For eight samples, the cushion comprised 5 mL PURE 2 M sucrose solution, 550 µL sucrose cushion buffer, 55 µL 0.1 M DTT, and 140 µL RNase inhibitor (5,600 U total; 700 U per sample). Filtered lysates were carefully overlaid onto 200 µL sucrose cushion and centrifuged at 16,100 × g for 45 min at 4 °C.

Following centrifugation, supernatants were removed and nuclei pellets were resuspended in 100 µL elution buffer (1× PBS, 1% fatty-acid-free BSA (MilliporeSigma, A1595), 2 mM EDTA, and RNase inhibitor, 40 U per 100 µL), transferred to fresh tubes to minimize debris carryover, and supplemented with an additional 100 µL elution buffer. Nuclei concentration and integrity were assessed using a LUNA automated cell counter (Logos Biosystems) with AO/PI staining (DeNovix, CD-AO-PI-1.5).

For FANS, nuclei were stained with DAPI (10 μL/mL, Roche 10236276001), with unstained controls included. Sorting was performed on a FACSAria Fusion (BD Biosciences) using a 70 µm nozzle, gating on DAPI-positive singlet nuclei. Sorted nuclei were collected into 1.5 ml LoBind tubes containing 10 µL elution buffer supplemented with 10 µL RNase inhibitor and kept on ice.

After nuclei were extracted, they were pelleted by centrifugation at 1,000 g for 5 min at 4 °C and resuspended with 100 μL 1x PBS with 0.1% w/v BSA and 1 U/μL RNase Inhibitor. Nuclei were then fixed by slowly adding 400 μL ice-cold methanol and stored at –20 °C overnight.

### Single-nucleus total RNA sequencing

Protocols for single-nucleus total RNA sequencing were based on TotalX protocols (*11*) and the single-nucleus total RNA sequencing protocols described in the STRS publication (*12*) with some modifications. Instead of the custom template-switching oligonucleotide (TSO) used in TotalX, a modified version of the 10x Genomics Single Cell 3’ v3.1 TSO containing only deoxyribonucleotides (5’-AAGCAGTGGTATCAACGCAGAGTACATGGG-3’, Integrated DNA Technologies) was used to prevent polyadenylation of the TSO. Uracil-DNA glycosylase treatment was omitted, and no custom spike-in primers were used for cDNA amplification. The second round of cDNA amplification was reduced to 5 cycles, and 20 ng of the resulting cDNA was further processed with the SEQuoia RiboDepletion Kit (Bio-Rad 17006487) following the manufacturer’s instructions. Library construction from the rRNA-depleted cDNA followed the standard 10x protocol with no modifications.

### Sequencing

All sequencing was performed by the CZBiohub San Francisco Genomics Platform. Completed sequencing libraries were loaded on Illumina NovaSeq 6000 S4 flow cells in sets of four libraries per lane or NovaSeq X 10B flow cells in sets of four libraries per two lanes with the goal of generating 50,000 to 75,000 reads per cell/nucleus.

### Sequencing data extraction

Sequencing data were de-multiplexed using bcl2fastq version 2.20.0.422. Adapter sequences and poly(A) regions were trimmed with Cutadapt (*85*) version 4.4, and rRNA reads were depleted *in silico* with bowtie2 (*86*) version 2.5.1. Resulting sequences were aligned to a custom reference constructed from the GRCh38.p14 primary assembly with gene annotations sourced from GENCODE (*87*) release 44, GtRNAdb (*88*) release 21, and miRbase release 22.1 (*89*), and quantified using STARsolo (*90*) version 2.7.11a with the following parameters: –-outFilterMismatchNoverLmax 0.05, –-outFilterMatchNmin 16, –-outFilterScoreMinOverLread 0, –-outFilterMatchNminOverLread 0, –-outFilterMultimapNmax50, –-soloMultiMappers EM.

### Data pre-processing and cell type annotations

Gene count tables were constructed from STARsolo outputs using Scanpy (*91*) version 1.10.1. For each library, ambient RNA reads were removed using CellBender (*92*) version 0.3.2, and doublets were filtered out with the implementation of Scrublet (*93*) in Scanpy. Cells and nuclei with fewer than 100 detected genes, fewer than 1,000 total UMIs, or greater than 20% mitochondrial UMIs were excluded. For each different tissue, samples were integrated with the implementation of Harmony (*94*) in Scanpy to generate unified visualizations and clusterings of cells and nuclei. Each cluster of cells and nuclei was then annotated initially using AnnDictionary (*13*) with claude-sonnet-4-20250514. Cell type labels were manually checked and corrected with known cell type marker genes.

### Integration with Tabula Sapiens 2.0

TSP33 whole-cell TotalX data were integrated with Tabula Sapiens 2.0 using the implementation of Harmony (*94*) in Scanpy (*91*) version 1.10.1, separately for each tissue and using only protein-coding genes. Cells from both datasets were then clustered using Scanpy’s implementation of the Leiden algorithm with default parameters.

### Differential gene expression analysis

All differential gene expression analysis (DGEA) was performed using Scanpy version 1.10.1. For all differentially expressed genes (DEGs), a log_2_ fold change > 1 and adjusted p-value < 0.05 for that cell type/modality compared to all others were required, with method=’wilcoxon’ and otherwise default parameters. To identify DEGs for each cell type, each modality was analyzed separately and then aggregated by taking the union of the DEGs from each modality.

### Evaluation of cell type specificity with the τ statistic

*τ* analysis was performed as previously described (*1*) with slight modifications. Annotated genes with fewer than 100 counts across the dataset were filtered out to avoid spurious alignments to low-confidence genes appearing as highly-specific genes. Additionally, only cell types with ≥100 cells and ≥100 nuclei were considered to limit bias due to dropout making genes appear more cell type-specific. The mean of the log-normalized expression for each gene was computed for each cell type. The *τ* statistic for each gene was then calculated as follows:

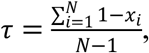

where *i* denotes the index of a particular cell type, *x*_*i*_ denotes the mean log-normalized expression in cell type *i*, and *N* denotes the total number of cell types considered.

All genes were ranked by *τ* in increasing order (least to most specific), and moving-window enrichment scores were calculated in 5,000 gene moving windows for each biotype by calculating the ratio of the proportion of genes of the biotype in the moving widow to the proportion of all filtered genes that belonged to that biotype.

### Evaluation of cell type specificity with expression entropy

To calculate the entropy specificity scores for a given gene, the mean expression values of that gene across all cell types were first normalized by dividing each cell type’s mean expression by the total mean expression across all cell types. The normalized Shannon entropy *H_norm_* was then calculated for that gene as follows:

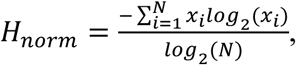

where *x_i_* represents the normalized mean expression in cell type *i* and *N* represents the number of cell types. The normalized entropy was then subtracted from one to obtain a metric ranging from 0 to 1 that increases with increasing specificity to mirror the specificity statistic *τ*.

### Identification of predominantly nuclear, predominantly cytoplasmic, and mixed genes

DEGs between whole cells and nuclei were identified for each cell type with at least 50 cells and 50 nuclei in the dataset. Genes with τ ≥ 0.85 were excluded to prevent genes with cell type-specific expression from appearing as genes with cell type-specific nuclear compartmentalization. Furthermore, only genes with adjusted p-value < 0.05 across at least half of cell types and absolute value of log_2_ fold change > 1 in at least one cell type were considered.

The resulting genes were split into three groups: “predominantly nuclear” if the log fold change > 0 (enriched in nuclei) for all cell types with adjusted p-value < 0.05, “predominantly cytoplasmic” if the log fold change < 0 (depleted in nuclei) for all cell types with adjusted p-value < 0.05, and “mixed” for genes that did not meet either of the previous criteria.

### Analysis for cell type specificity of nuclear compartmentalization profiles

To evaluate cell type specificity of nuclear compartmentalization, a trinarized matrix representing nuclear enrichment status for each gene in each tissue-cell type combination was constructed: +1 if log fold change > 1 and adjusted p-value < 0.05 for that gene in that tissue-cell type, –1 if log fold change < –1 and adjusted p-value < 0.05, and 0 otherwise. Pairwise cosine similarities were then computed between all tissue-cell type combinations to generate a similarity matrix, which was then aggregated to the cell type level by averaging across tissues for each cell type. For within-cell type comparisons, only pairs of distinct tissues were included to avoid inflating similarity estimates through self-correlations of the same tissue-cell type. The mean of the within-cell type comparisons was then recomputed for 1,000 permutations of tissue-cell type labels to generate the null distribution for the permutation test.

### Analysis for cell type specificity of tRNA repertoires

Only the whole-cell data were considered for all analyses of tRNAs. Furthermore, tRX and undetermined tRNA genes were removed, and tRNA gene variants were aggregated by summing the counts for each tRNA isoacceptor family. Only tissue-cell types with at least 500 tRNA counts total were considered. The tRNA counts for all cells in each tissue-cell type were then summed, normalized to 10,000 counts per tissue-cell type, log transformed, and scaled separately for each sample. Pairwise Pearson correlations were then computed for all tissue-cell type combinations. For each pair of cell types that were found in at least three different tissues, mean correlations were calculated across all tissue combinations, with within-cell type correlations excluding self-comparisons. The means of each set of within-cell type comparisons were then recomputed for 1,000 permutations of tissue-cell type labels to generate null distributions for each cell type.

### Evaluation of theoretical translation efficiency

Theoretical translation efficiency (tTE) was previously defined as the Spearman’s correlation coefficient between the tRNA isotype abundance profile and the amino acid usage profile of each cell type (*64*). To compute tRNA isotype abundance profiles, tRNA expression values were aggregated at the amino acid level for each cell, normalized to 10,000 counts, and averaged for each cell type. Amino acid usage for each gene was derived from the protein-coding transcript translation sequences in GENCODE release 44. For genes with multiple isoforms, the mean amino acid composition across all annotated isoforms was used. The mean amino acid usage profile for each cell type was computed by summing amino acid counts from all detected protein-coding genes in each cell, normalized to 10,000 counts for each cell, and averaging for each cell type. Finally, tTE for each cell type was computed as the Spearman’s correlation between its mean normalized tRNA abundance and amino acid usage profiles. The tTEs were then recomputed for 1,000 permutations of amino acid labels or of cell type labels to generate null distributions.

### Assignment of cell-cycle stages

For cell types that are known to divide, cell-cycle phases were assigned using the score_genes_cell_cycle function of Scanpy (*91*) version 1.10.1 with a standard list of protein-coding marker genes for the S and G2/M phases (*66*).

### Analysis of gene clusters across cell-cycle stages

Following the analysis described in the smart-seq-total publication (*95*), the top 250 differentially expressed protein coding genes for each cell-cycle phase and all non-coding genes with average log-normalized expression > 0.05 in at least one phase across all cells were used for the clustering analyses for cells and nuclei separately. For each gene that met these criteria, the mean expression in each phase for each cell type was calculated. The resulting cell type-gene by phase matrix was then z-scored across phases, L2-normalized, and clustered into twelve groups with K-means clustering. Groups were manually combined into six groups based on high/low expression in a given phase: G1+ (high in G1), G1– (low in G1), S+, S-, G2/M+, and G2/M-. Genes were deemed consistently assigned if a single combined group accounted for ≥60% of their detected cell types and was represented in at least one-third of the cell types considered.

### Analysis of senescence-associated ncRNA genes

To analyze senescence-associated ncRNAs, cells expressing *CDKN2A* but not expressing *MKI67* were identified and used as putative senescent cells. To uncover senescence-associated changes in ncRNA expression patterns, biotype abundances were calculated for *CDKN2A*+ *MKI67*− cells and compared with other cells across cell types. To identify senescence-associated ncRNAs, DGEA was performed between *CDKN2A*+ *MKI67*− cells and the remaining cells within the same cell type. Only genes with a log fold change > 0.5, adjusted p-values < 0.01, and expression in >50% of *CDKN2A*+ *MKI67*− cells were considered enriched in putative senescent cells.

## Supplementary Figures

**Supplementary Figure 1.**
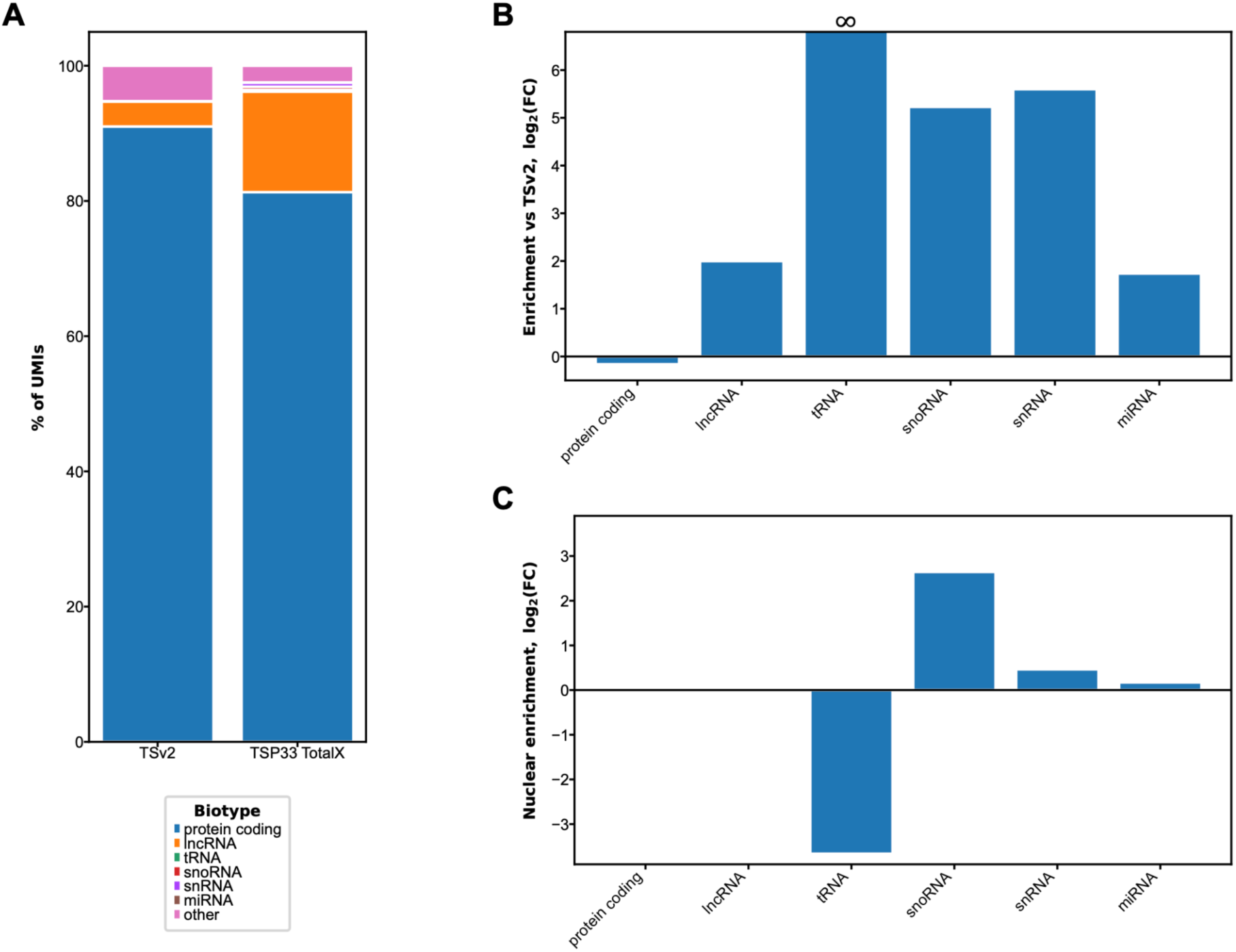
ncRNA enrichment. A) Stacked bar chart showing proportions of UMIs detected belonging to selected biotypes in Tabula Sapiens 2.0 (TSv2, left) and the TSP33 TotalX dataset (right). B) Bar chart showing the log_2_ fold change in UMI proportions for selected biotypes in the total RNA sequencing dataset compared to TSv2. No tRNAs were reported in TSv2, as denoted by the ∞ above the tRNA bar. C) Bar chart showing the log_2_ fold change of UMI proportions for selected biotypes in the single-nucleus portion of the total RNA sequencing dataset compared to the single-cell portion.

**Supplementary Figure 2.**
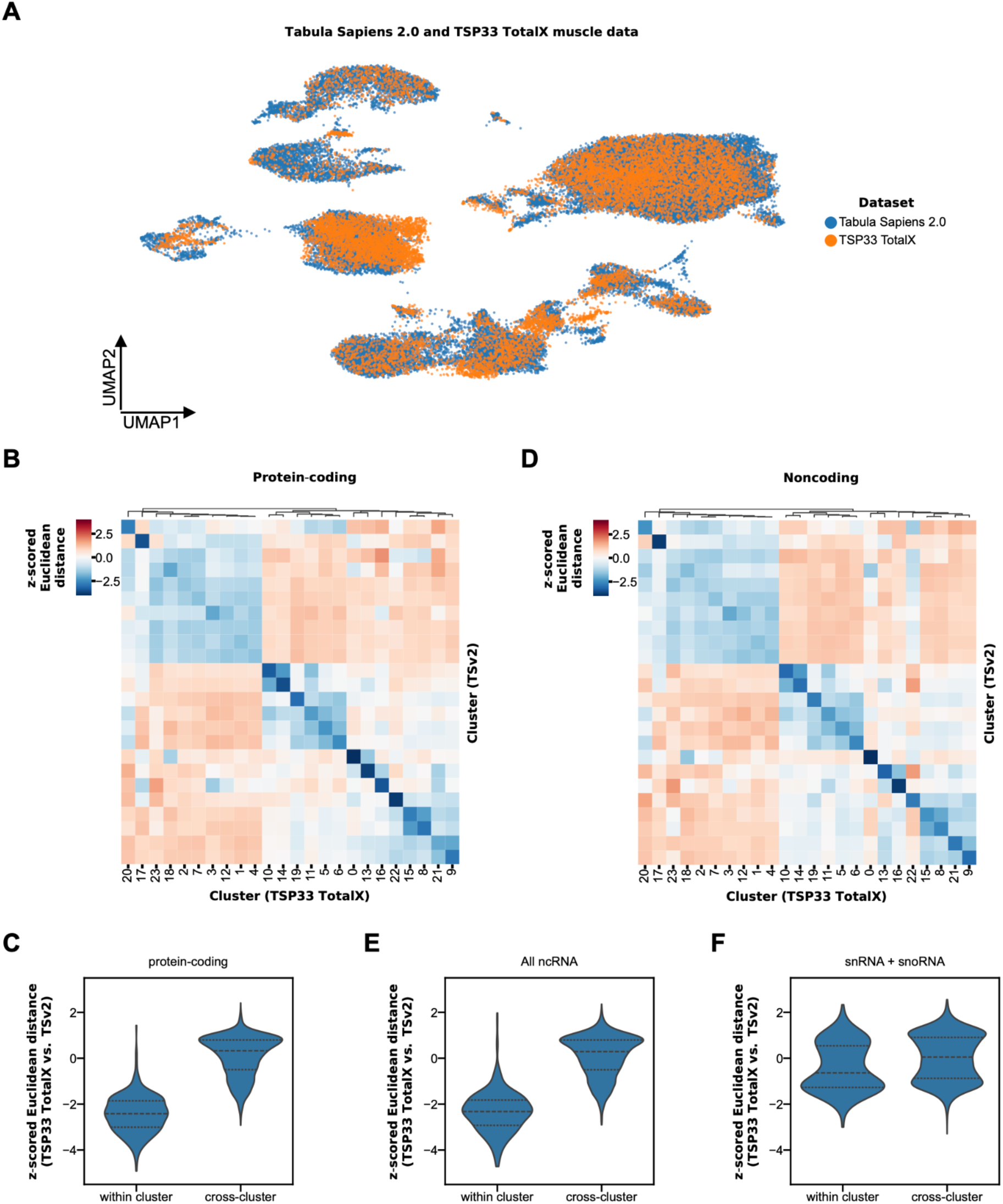
Concordance with Tabula Sapiens 2.0. A) Representative Uniform Manifold Approximation and Projection (UMAP) visualization of Tabula Sapiens 2.0 and TSP33 whole-cell TotalX data after integration for muscle, colored by dataset. B) Representative heat map showing z-scored Euclidean distances between cluster-mean protein-coding expression profiles from Tabula Sapiens 2.0 muscle (rows) and from TSP33 TotalX muscle (columns). Columns are hierarchically clustered by Euclidean distance, as indicated by the dendrogram above. Rows are displayed in the same order as the columns to align corresponding clusters, such that diagonal entries represent comparisons of Tabula Sapiens 2.0 to TSP33 TotalX within groups of similar cells. C) Violin plots of z-scored Euclidean distances between cluster-mean protein-coding expression profiles from Tabula Sapiens 2.0 and TSP33 TotalX across all tissues shared between datasets, with matched (within cluster) and mismatched (cross-cluster) cluster pairs. D) Representative heat map showing z-scored Euclidean distances between cluster-mean non-coding expression profiles from Tabula Sapiens 2.0 muscle (rows) and from TSP33 TotalX muscle (columns). Columns are hierarchically clustered by Euclidean distance, as indicated by the dendrogram above. Rows are displayed in the same order as the columns. E) Violin plots of z-scored Euclidean distances between cluster-mean non-coding expression profiles from Tabula Sapiens 2.0 and TSP33 TotalX across all tissues shared between datasets, with matched (within cluster) and mismatched (cross-cluster) cluster pairs. F) Violin plots of z-scored Euclidean distances between cluster-mean snoRNA and snRNA expression profiles from Tabula Sapiens 2.0 and TSP33 TotalX across all tissues shared between datasets, with matched (within cluster) and mismatched (cross-cluster) cluster pairs.

**Supplementary Figure 3.**
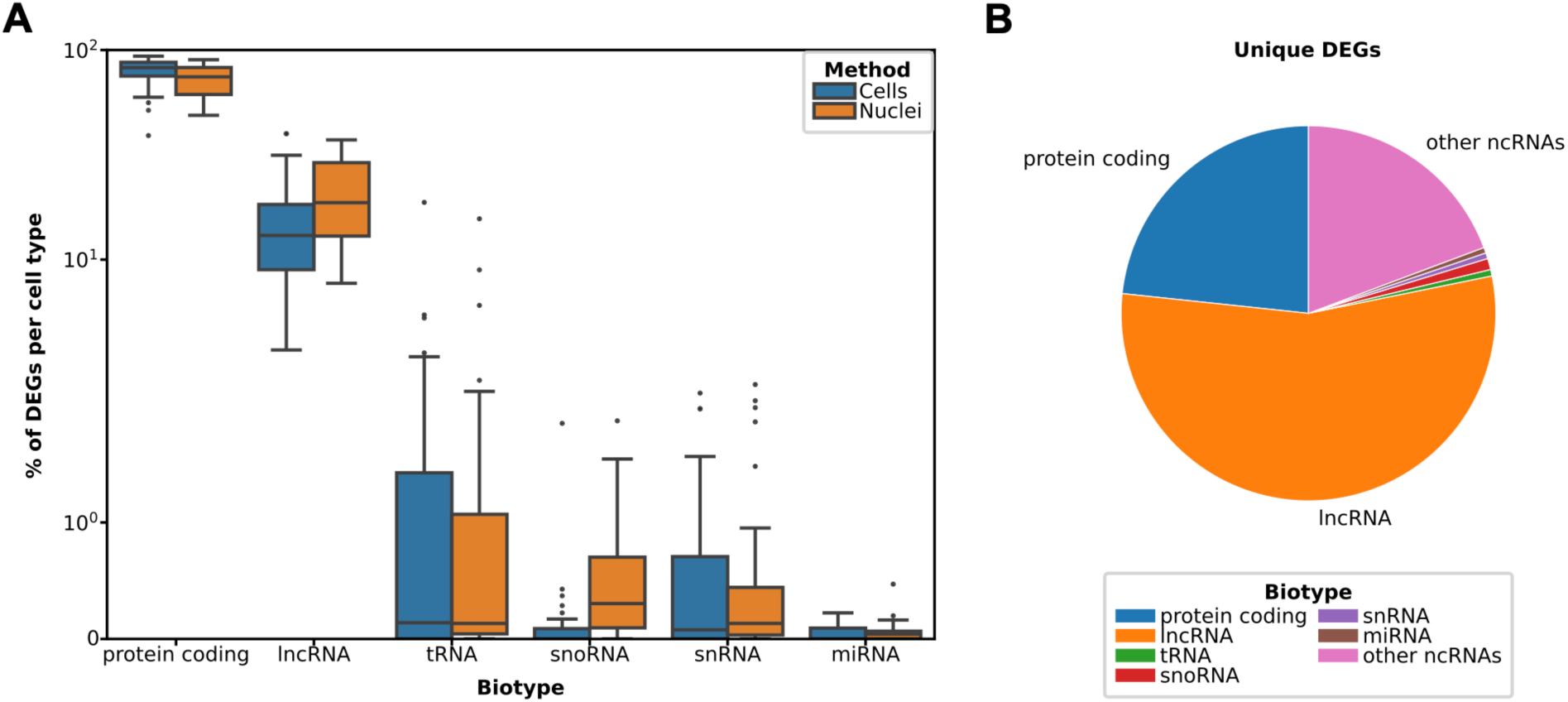
Shared and unique differentially expressed genes between cell types. A) Box plots showing the distributions of proportions of differentially expressed (DEGs) belonging to selected biotypes, colored by method. B) Pie chart showing the biotype composition of DEGs uniquely identified in a single cell type (“unique DEGs”).

**Supplementary Figure 4.**
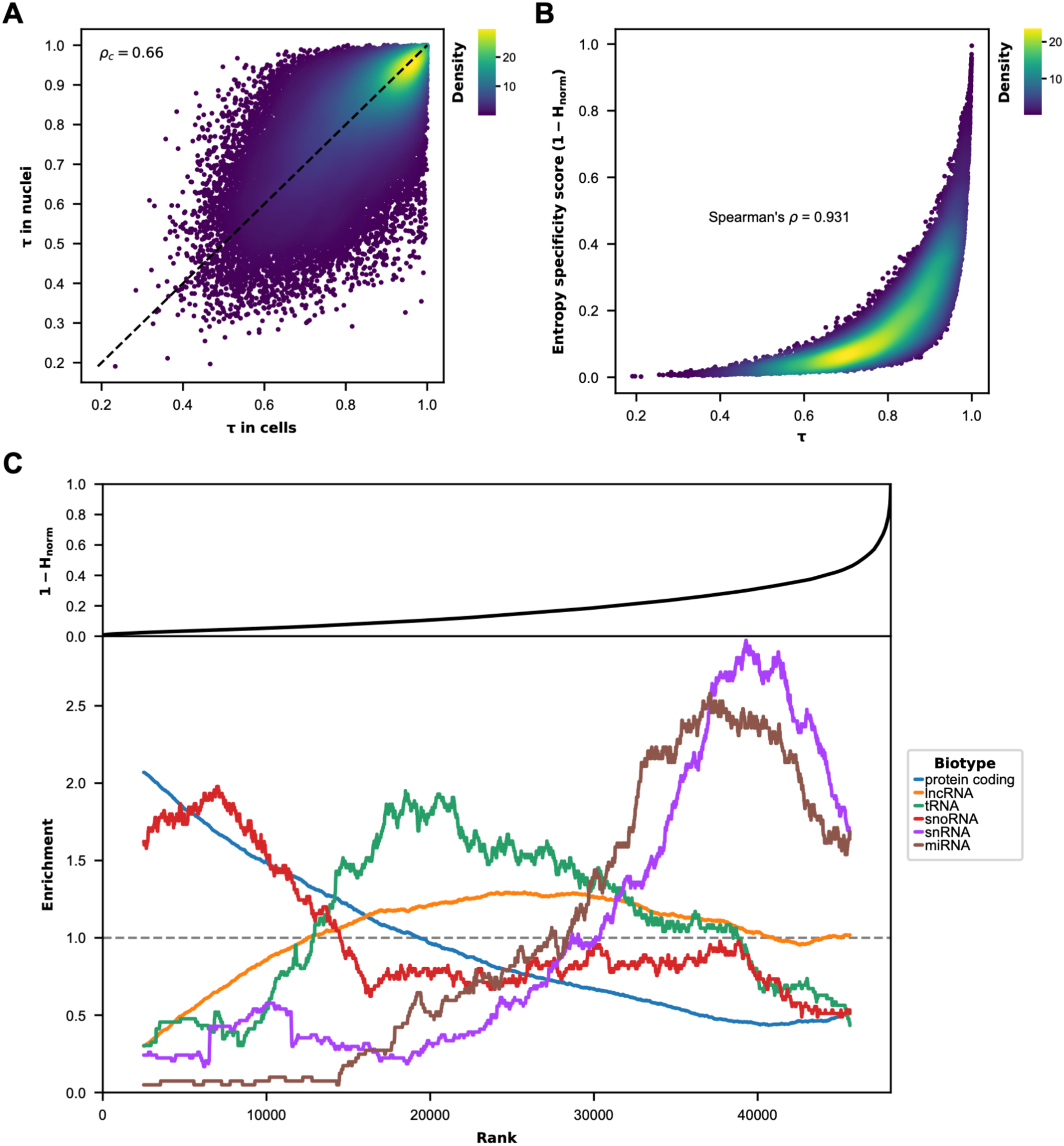
Comparison of cell type specificity metrics. A) Scatter plot comparing the specificity statistic τ calculated from the single-nucleus portion of the total RNA sequencing dataset to that calculated from the single-cell portion for each gene, colored by density of points. The one-to-one line (black dashed line) and concordance correlation coefficient (*p_c_*) are shown. B) Scatter plot comparing expression entropy to specificity index *τ* for each gene. C) Top: expression entropy *τ* plotted against gene rank, with genes ordered by decreasing entropy. Bottom: enrichment of each biotype within a moving window across the same gene ranking, normalized by the total number of genes of that biotype. An enrichment value of 1 indicates that the local proportion of genes from a given biotype equals its overall proportion across all annotated genes with ≥ 50 counts across the dataset.

**Supplementary Figure 5.**
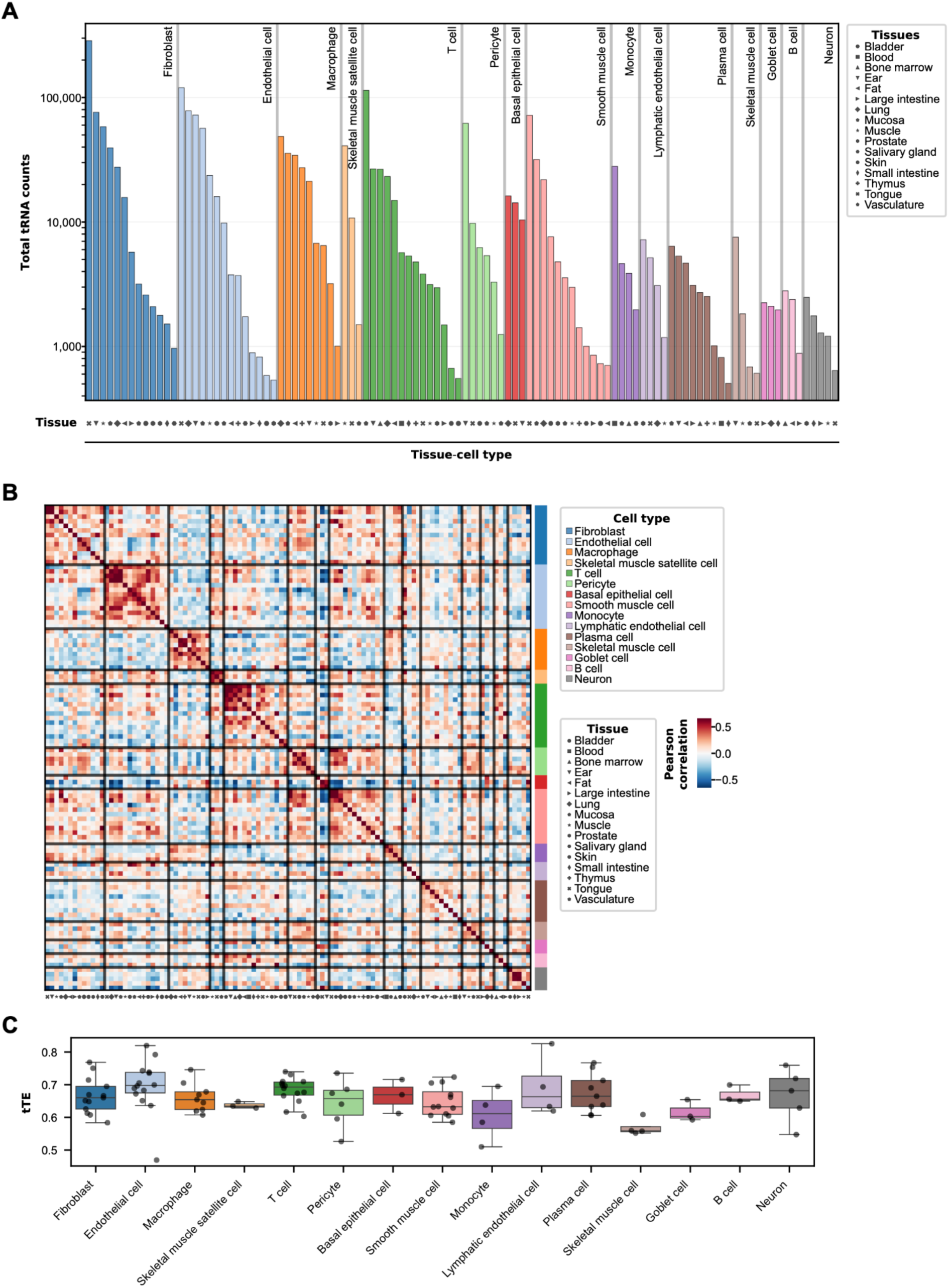
tRNA profiles by tissue-cell type. A) Bar charts showing the total number of tRNA UMIs detected in each tissue-cell type combination. Bars are grouped and colored by cell type, and tissues for each cell type are ordered in descending number of UMIs. B) Heat map of Pearson correlation coefficients between tRNA expression profiles for tissue-cell type combinations, grouped by cell type (color bar, right) and ordered by tissue (symbols, bottom) as in panel A. Black lines delineate cell type group boundaries. C) Box plots of theoretical translation efficiency (tTE) values for each cell type, with points indicating individual tissue-cell type combinations within each cell type.

**Supplementary Figure 6.**
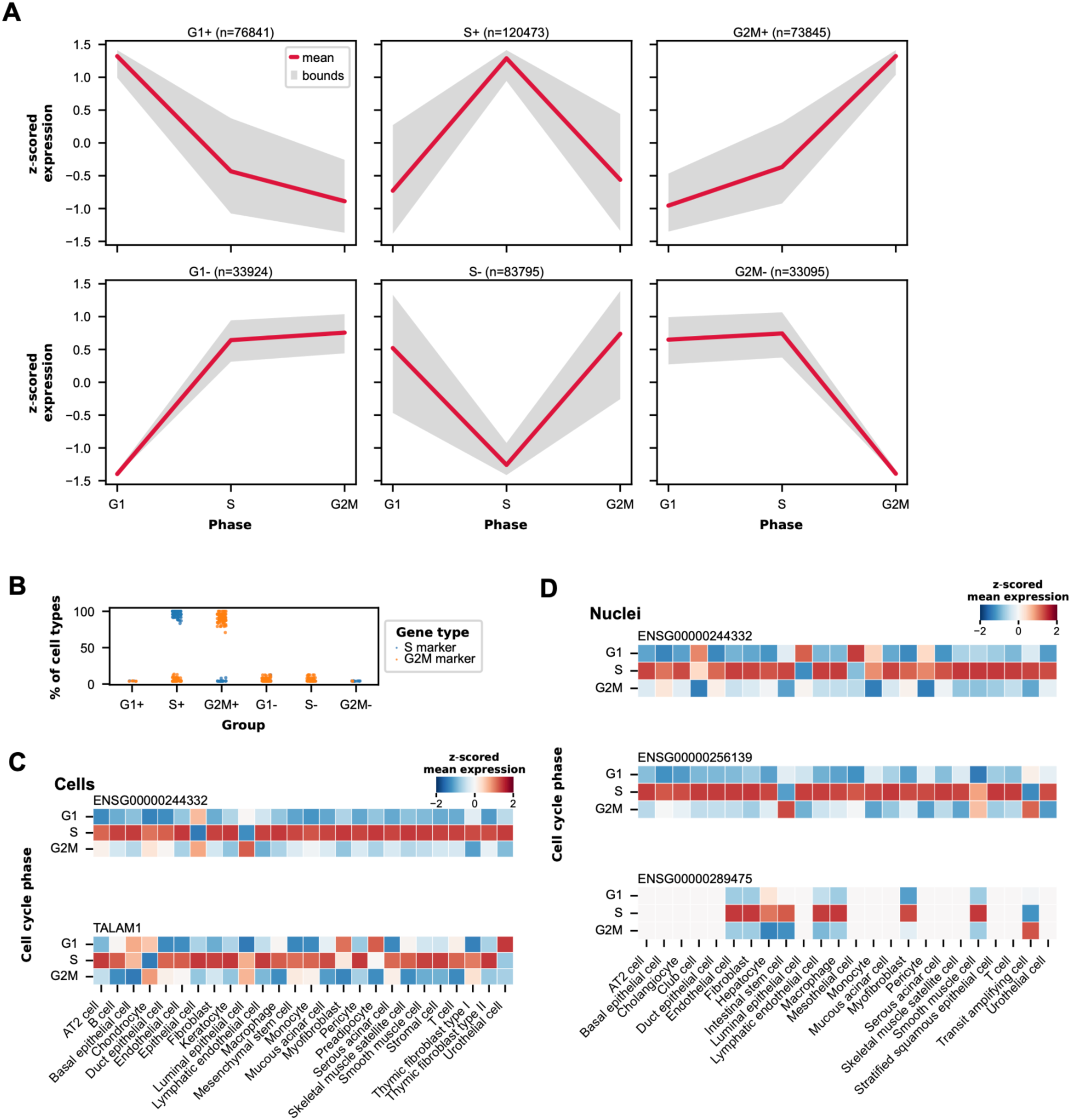
Expression profiles across the cell cycle. A) Expression profiles of cell cycle-dependent gene clusters. For each cell type that is known to divide, the mean expression values of selected protein-coding and ncRNA genes were calculated across cell-cycle phases (G1, S, and G2/M). The resulting cell type-gene expression profiles were grouped into six clusters based on high (+) or low (−) expression in each phase. For each cluster, the central red line denotes the mean expression profile across phases, and the gray shaded region spans the minimum and maximum values in each phase. B) Scatter plot showing, for each protein-coding gene used as a marker for S (blue) or G2/M (orange) phase, the proportion of cell types in which that gene is assigned to each cluster. C) Heat maps showing Z-scored mean expression values across cell type and cell-cycle phase for ncRNA genes that were consistently upregulated in S phase in the single-cell portion of the total RNA sequencing dataset. D) Heat maps showing Z-scored mean expression values across cell type and cell-cycle phase for ncRNA genes that were consistently upregulated in S phase in the single-nucleus portion of the total RNA sequencing dataset.

## The Tabula Sapiens Consortium

Stephen R. Quake^1,2^

Robert C Jones^1^

Mark Krasnow^3,4^

Angela Oliveira Pisco^5^

Julia Salzman^3,6^

Nir Yosef^5,7,8^

George Crowley^1^

Siyu He^1,6^

Madhav Mantri^1^

Jessie Aguirre^9^

Ron Garner^9^

Sal Guerrero^9^

William Harper^9^

Resham Irfan^9^

Sophia Mahfouz^9^

Ravi Ponnusamy^9^

Bhavani A. Sanagavarapu^9^

Ahmad Salehi^9^

Ivan Sampson^9^

Chloe Tang^9^

Alan G. Cheng^10^

James M. Gardner^11,12^

Burnett Kelly^9,13^

Thurman Slone^9^

Zifa Wang^9^

Anika Choudhury^1,5^

Sheela Crasta^1^

Chen Dong^1,5^

Marcus L. Forst^1^

Douglas E. Henze^1^

Jaeyoon Lee^1^

Maurizio Morri^5^

Serena Y. Tan^14^

Sevahn K. Vorperian^15,16^

Lynn Yang^1,5^

Marcela Alcántara-Hernádez^17^

Julian Berg^18^

Dhruv Bhatt^19^

Sara Billings^10^

Andrès Gottfried-Blackmore^17,20^

Jamie Bozeman^19^

Simon Bucher^21^

Elisa Caffrey^22^

Amber Casillas^23^

Rebecca Chen^22^

Matthew Choi^21^

Rebecca N. Culver^24^

Ivana Cvijovic^1,2^

Ke Ding^25^

Hala Shakib Dhowre^26^

Hua Dong^25^

Kenneth Donaville^21^

Lauren Duan^18^

Xiaochen Fan^19^

Mariko H. Foecke^23^

Francisco X. Galdos^18^

Eliza A. Gaylord^23^

Karen Gonzales^19^

William R. Goodyer^27^

Michelle Griffin^28^

Yuchao Gu^3,29,30^

Shuo Han^25^

Jun Yan He^22^

Paul Heinrich^18^

Rebeca Arroyo Hornero^17^

Keliana Hui^22^

Juan C. Irwin^23^

SoRi Jang^3^

Annie Jensen^18,31^

Saswati Karmakar^14,24^

Jengmin Kang^32^

Hailey Kang^21^

Soochi Kim^32^

Stewart J. Kim^3,29,30^

William Kong^25^

Mallory A. Laboulaye^25^

Daniel Lee^18^

Gyehyun Lee^33^

Elise Lelou^21^

Anping Li^25^

Baoxiang Li^26^

Wan-Jin Lu^25^

Hayley Raquer-McKay^17^

Elvira Mennillo^33^

Lindsay Moore^34^

Elena Montauti^22^

Karim Mrouj^25^

Shravani Mukherjee^26^

Patrick Neuhöfer^3,29,30^

Saphia Nguyen^21^

Honor Paine^21^

Jennifer B. Parker^25,28^

Julia Pham^22^

Kiet T. Phong^35^

Pratima Prabala^19^

Zhen Qi^25^

Joshua Quintanilla^18,31^

Iulia Rusu^33^

Ali Reza Rais Sadati^18^

Bronwyn Scott^26^

David Seong^17^

Hosu Sin^36^

Hanbing Song^37^

Bikem Soyur^23^

Sean Spencer^17,20^

Varun R. Subramaniam^26^

Michael Swift^1^

Aditi Swarup^26^

Greg Szot^11,12^

Aris Taychameekiatchai^21^

Emily Trimm^19^

Stefan Veizades^18,31^

Sivakamasundari Vijayakumar^25^

Kim Chi Vo^23^

Tian Wang^34^

Timothy Wu^3^

Yinghua Xie^18,31^

William Yue^21^

Zue Zhang^3^

Angela Detweiler^5^

Honey Mekonen^5^

Norma F. Neff^5^

Sheryl Paul^5^

Amanda Seng^5^

Jia Yan^5^

Deana Rae Crystal Colburg^38^

Balint Laszlo Forgo^14^

Luca Ghita^17^

Frank McCarthy^39^

Aditi Agrawal^5^

Alina Isakova^1^

Kavita Murthy^1^

Alexandra Psaltis^1^

Wenfei Sun^1^

Kyle Awayan^5^

Pierre Boyeau^40^

Robrecht Cannoodt^41–43^

Leah Dorman^5^

Samuel D’Souza^5^

Can Ergen^7,40^

Justin Hong^44^

Harper Hua^6^

Erin McGeever^5^

Antoine de Morree^32,45,46^

Luise A. Seeker^1^

Alexander J. Tarashansky^5^

Astrid Gillich^3^

Taha A. Jan^47^

Angela Ling^47^

Abhishek Murti^21^

Nikita Sajai^21^

Ryan M. Samuel^48^

Juliane Winkler^49,50^

Steven E. Artandi^3,29,30^

Philip A. Beachy^25,31,51^

Mike F. Clarke^25^

Zev Gartner^5,52^

Linda C. Giudice^53^

Franklin W. Huang^37,54^

Juliana Idoyaga^17,55^

Michael G. Kattah^33^

Christin S. Kuo^56^

Diana J. Laird^23^

Michael T. Longaker^25,57^

Patricia Nguyen^18,31,58^

David Y. Oh^22^

Thomas A. Rando^32^

Kristy Red-Horse^19^

Bruce Wang^21^

Albert Y. Wu^26^

Sean M. Wu^18,31^

Bo Yu^36^

James Zou^6,59^

1. Department of Bioengineering, Stanford University; Stanford, CA, USA.
2. Department of Applied Physics, Stanford University, Stanford, CA, USA.
3. Department of Biochemistry, Stanford University School of Medicine, Stanford, CA, USA.
4. Howard Hughes Medical Institute, USA.
5. Chan Zuckerberg Biohub, San Francisco, CA, USA.
6. Department of Biomedical Data Science, Stanford University, Stanford, CA, USA.
7. Center for Computational Biology, University of California Berkeley, Berkeley, CA, USA.
8. Ragon Institute of MGH, MIT and Harvard, Cambridge, MA, USA.
9. Donor Network West, San Ramon, CA, USA.
10. Department of Otolaryngology-Head and Neck Surgery, Stanford University School of Medicine, Stanford, California, USA.
11. Department of Surgery, University of California San Francisco, San Francisco, CA, USA.
12. Diabetes Center, University of California San Francisco, San Francisco, CA, USA.
13. DCI Donor Services, Sacramento, CA, USA.
14. Department of Pathology, Stanford University School of Medicine, Stanford, CA, USA.
15. Department of Chemical Engineering, Stanford University, Stanford, CA, USA.
16. Sarafan ChEM-H, Stanford University, Stanford, CA, USA.
17. Department of Microbiology and Immunology, Stanford University School of Medicine, Stanford, CA, USA.
18. Stanford Cardiovascular Institute, Stanford CA, USA.
19. Department of Biology, Stanford University, Stanford, CA, USA.
20. Division of Gastroenterology, Department of Medicine, Stanford University School of Medicine, Stanford, CA, USA.
21. Department of Medicine and Liver Center, University of California San Francisco, San Francisco, CA, USA.
22. Division of Hematology/Oncology, Department of Medicine, University of California San Francisco, San Francisco, CA, USA.
23. Department of Ob/Gyn and Reproductive Sciences, Eli and Edythe Broad Center for Regeneration Medicine and Stem Cell Research, University of California, San Francisco.
24. Department of Genetics, Stanford University School of Medicine, Stanford, CA, USA.
25. Institute for Stem Cell Biology and Regenerative Medicine, Stanford University School of Medicine, Stanford, CA, USA.
26. Department of Ophthalmology, Stanford University School of Medicine, Stanford, CA, USA.
27. Department of Pediatrics, Division of Cardiology, Stanford University School of Medicine, Stanford, CA, USA.
28. Department of Surgery, Division of Plastic and Reconstructive Surgery, Stanford University School of Medicine, Stanford, CA, USA.
29. Stanford Cancer Institute, Stanford University School of Medicine, Stanford, CA, USA.
30. Department of Medicine, Division of Hematology, Stanford University School of Medicine, Stanford, CA, USA.
31. Department of Medicine, Division of Cardiovascular Medicine, Stanford University, Stanford, CA, USA.
32. Department of Neurology and Neurological Sciences, Stanford University School of Medicine, Stanford, CA, USA.
33. Division of Gastroenterology, Department of Medicine, University of California, San Francisco, San Francisco, CA, USA.
34. Division of Pediatric Otolaryngology Stanford University School of Medicine, Stanford, CA, USA.
35. Department of Bioengineering and Therapeutic Sciences, University of California, San Francisco, San Francisco, CA, USA.
36. Department of OB/GYN Stanford University, Palo Alto, CA, USA.
37. Division of Hematology and Oncology, Department of Medicine, Bakar Computational Health Sciences Institute, Institute for Human Genetics, University of California San Francisco, San Francisco, CA, USA.
38. Stanford Health Care, Stanford CA, USA.
39. Mass Spectrometry Platform, Chan Zuckerberg Biohub, Stanford, CA, USA.
40. Department of Electrical Engineering and Computer Sciences, University of California Berkeley, Berkeley, CA, USA.
41. Data Intuitive, Flanders, Belgium.
42. Data Mining and Modelling for Biomedicine group, VIB Center for Inflammation Research, Ghent, Belgium.
43. Department of Applied Mathematics, Computer Science, and Statistics, Ghent University, Ghent, Belgium.
44. Department of Computer Science, Columbia University; New York, NY, USA.
45. Department of Biomedicine, Aarhus University, Aarhus, Denmark.
46. Paul F. Glenn Center for the Biology of Aging, Stanford University School of Medicine, Stanford, CA, USA.
47. Department of Otolaryngology, Vanderbilt University Medical Center, Nashville, TN, USA.
48. Department of Cellular Molecular Pharmacology, University of California, San Francisco, San Francisco, CA, USA.
49. Department of Cell & Tissue Biology, University of California San Francisco, San Francisco, CA, USA.
50. Center for Cancer Research Medical University of Vienna Borschkegasse 8a 1090 Vienna, Austria.
51. Department of Urology, Stanford University School of Medicine, Stanford, CA, USA.
52. Department of Pharmaceutical Chemistry, University of California San Francisco, San Francisco, CA.
53. Center for Reproductive Sciences, Department of Obstetrics, Gynecology and Reproductive Sciences, University of California San Francisco, San Francisco, CA, USA.
54. Division of Hematology/Oncology, Department of Medicine, San Francisco Veterans Affairs Health Care System, San Francisco, CA, USA.
55. Pharmacology and Molecular Biology Departments, Schools of Medicine and Biological Sciences, University of California, San Diego, CA, USA.
56. Department of Pediatrics, Division of Pulmonary Medicine, Stanford University, Stanford, CA, USA.
57. Department of Surgery, Stanford University School of Medicine, Stanford, CA, USA.
58. Veterans Affairs Palo Alto Health Care System, Palo Alto, CA, USA.
59. Department of Computer Science, Stanford University, Stanford, CA.

## Notes

### Competing Interest Statement

The authors have declared no competing interest.

## References

1. T. T. S. Consortium, S. R. Quake, Tabula Sapiens reveals transcription factor expression, senescence effects, and sex-specific features in cell types from 28 human organs and tissues. bioRxiv [Preprint] (2025). 10.1101/2024.12.03.626516.

2. THE TABULA SAPIENS CONSORTIUM, The Tabula Sapiens: A multiple-organ, single-cell transcriptomic atlas of humans. Science 376, eabl4896 (2022).

3. C. Domínguez Conde, C. Xu, L. B. Jarvis, D. B. Rainbow, S. B. Wells, T. Gomes, S. K. Howlett, O. Suchanek, K. Polanski, H. W. King, L. Mamanova, N. Huang, P. A. Szabo, L. Richardson, L. Bolt, E. S. Fasouli, K. T. Mahbubani, M. Prete, L. Tuck, N. Richoz, Z. K. Tuong, L. Campos, H. S. Mousa, E. J. Needham, S. Pritchard, T. Li, R. Elmentaite, J. Park, E. Rahmani, D. Chen, D. K. Menon, O. A. Bayraktar, L. K. James, K. B. Meyer, N. Yosef, M. R. Clatworthy, P. A. Sims, D. L. Farber, K. Saeb-Parsy, J. L. Jones, S. A. Teichmann, Cross-tissue immune cell analysis reveals tissue-specific features in humans. Science 376, eabl5197 (2022).

4. L. Sikkema, C. Ramírez-Suástegui, D. C. Strobl, T. E. Gillett, L. Zappia, E. Madissoon, N. S. Markov, L.-E. Zaragosi, Y. Ji, M. Ansari, M.-J. Arguel, L. Apperloo, M. Banchero, C. Bécavin, M. Berg, E. Chichelnitskiy, M.-I. Chung, A. Collin, A. C. A. Gay, J. Gote-Schniering, B. Hooshiar Kashani, K. Inecik, M. Jain, T. S. Kapellos, T. M. Kole, S. Leroy, C. H. Mayr, A. J. Oliver, M. von Papen, L. Peter, C. J. Taylor, T. Walzthoeni, C. Xu, L. T. Bui, C. De Donno, L. Dony, A. Faiz, M. Guo, A. J. Gutierrez, L. Heumos, N. Huang, I. L. Ibarra, N. D. Jackson, P. Kadur Lakshminarasimha Murthy, M. Lotfollahi, T. Tabib, C. Talavera-López, K. J. Travaglini, A. Wilbrey-Clark, K. B. Worlock, M. Yoshida, Lung Biological Network Consortium, M. van den Berge, Y. Bossé, T. J. Desai, O. Eickelberg, N. Kaminski, M. A. Krasnow, R. Lafyatis, M. Z. Nikolic, J. E. Powell, J. Rajagopal, M. Rojas, O. Rozenblatt-Rosen, M. A. Seibold, D. Sheppard, D. P. Shepherd, D. D. Sin, W. Timens, A. M. Tsankov, J. Whitsett, Y. Xu, N. E. Banovich, P. Barbry, T. E. Duong, C. S. Falk, K. B. Meyer, J. A. Kropski, D. Pe’er, H. B. Schiller, P. R. Tata, J. L. Schultze, S. A. Teichmann, A. V. Misharin, M. C. Nawijn, M. D. Luecken, F. J. Theis, An integrated cell atlas of the lung in health and disease. Nat. Med. 29, 1563–1577 (2023).

5. T. Kumar, K. Nee, R. Wei, S. He, Q. H. Nguyen, S. Bai, K. Blake, M. Pein, Y. Gong, E. Sei, M. Hu, A. K. Casasent, A. Thennavan, J. Li, T. Tran, K. Chen, B. Nilges, N. Kashikar, O. Braubach, B. Ben Cheikh, N. Nikulina, H. Chen, M. Teshome, B. Menegaz, H. Javaid, C. Nagi, J. Montalvan, T. Lev, S. Mallya, D. F. Tifrea, R. Edwards, E. Lin, R. Parajuli, S. Hanson, S. Winocour, A. Thompson, B. Lim, D. A. Lawson, K. Kessenbrock, N. Navin, A spatially resolved single-cell genomic atlas of the adult human breast. Nature 620, 181–191 (2023).

6. J. C. Melms, J. Biermann, H. Huang, Y. Wang, A. Nair, S. Tagore, I. Katsyv, A. F. Rendeiro, A. D. Amin, D. Schapiro, C. J. Frangieh, A. M. Luoma, A. Filliol, Y. Fang, H. Ravichandran, M. G. Clausi, G. A. Alba, M. Rogava, S. W. Chen, P. Ho, D. T. Montoro, A. E. Kornberg, A. S. Han, M. F. Bakhoum, N. Anandasabapathy, M. Suárez-Fariñas, S. F. Bakhoum, Y. Bram, A. Borczuk, X. V. Guo, J. H. Lefkowitch, C. Marboe, S. M. Lagana, A. Del Portillo, E. J. Tsai, E. Zorn, G. S. Markowitz, R. F. Schwabe, R. E. Schwartz, O. Elemento, A. Saqi, H. Hibshoosh, J. Que, B. Izar, A molecular single-cell lung atlas of lethal COVID-19. Nature 595, 114–119 (2021).

7. A.-C. Villani, R. Satija, G. Reynolds, S. Sarkizova, K. Shekhar, J. Fletcher, M. Griesbeck, A. Butler, S. Zheng, S. Lazo, L. Jardine, D. Dixon, E. Stephenson, E. Nilsson, I. Grundberg, D. McDonald, A. Filby, W. Li, P. L. De Jager, O. Rozenblatt-Rosen, A. A. Lane, M. Haniffa, A. Regev, N. Hacohen, Single-cell RNA-seq reveals new types of human blood dendritic cells, monocytes, and progenitors. Science 356, eaah4573 (2017).

8. I. Dunham, A. Kundaje, S. F. Aldred, P. J. Collins, C. A. Davis, F. Doyle, C. B. Epstein, S. Frietze, J. Harrow, R. Kaul, J. Khatun, B. R. Lajoie, S. G. Landt, B.-K. Lee, F. Pauli, K. R. Rosenbloom, P. Sabo, A. Safi, A. Sanyal, N. Shoresh, J. M. Simon, L. Song, N. D. Trinklein, R. C. Altshuler, E. Birney, J. B. Brown, C. Cheng, S. Djebali, X. Dong, I. Dunham, J. Ernst, T. S. Furey, M. Gerstein, B. Giardine, M. Greven, R. C. Hardison, R. S. Harris, J. Herrero, M. M. Hoffman, S. Iyer, M. Kellis, J. Khatun, P. Kheradpour, A. Kundaje, T. Lassmann, Q. Li, X. Lin, G. K. Marinov, A. Merkel, A. Mortazavi, S. C. J. Parker, T. E. Reddy, J. Rozowsky, F. Schlesinger, R. E. Thurman, J. Wang, L. D. Ward, T. W. Whitfield, S. P. Wilder, W. Wu, H. S. Xi, K. Y. Yip, J. Zhuang, B. E. Bernstein, E. Birney, I. Dunham, E. D. Green, C. Gunter, M. Snyder, M. J. Pazin, R. F. Lowdon, L. A. L. Dillon, L. B. Adams, C. J. Kelly, J. Zhang, J. R. Wexler, E. D. Green, P. J. Good, E. A. Feingold, B. E. Bernstein, E. Birney, G. E. Crawford, J. Dekker, L. Elnitski, P. J. Farnham, M. Gerstein, M. C. Giddings, T. R. Gingeras, E. D. Green, R. Guigó, R. C. Hardison, T. J. Hubbard, M. Kellis, W. J. Kent, J. D. Lieb, E. H. Margulies, R. M. Myers, M. Snyder, J. A. Stamatoyannopoulos, S. A. Tenenbaum, Z. Weng, K. P. White, B. Wold, J. Khatun, Y. Yu, J. Wrobel, B. A. Risk, H. P. Gunawardena, H. C. Kuiper, C. W. Maier, L. Xie, X. Chen, M. C. Giddings, B. E. Bernstein, C. B. Epstein, N. Shoresh, J. Ernst, P. Kheradpour, T. S. Mikkelsen, S. Gillespie, A. Goren, O. Ram, X. Zhang, L. Wang, R. Issner, M. J. Coyne, T. Durham, M. Ku, T. Truong, L. D. Ward, R. C. Altshuler, M. L. Eaton, M. Kellis, S. Djebali, C. A. Davis, A. Merkel, A. Dobin, T. Lassmann, A. Mortazavi, A. Tanzer, J. Lagarde, W. Lin, F. Schlesinger, C. Xue, G. K. Marinov, J. Khatun, B. A. Williams, C. Zaleski, J. Rozowsky, M. Röder, F. Kokocinski, R. F. Abdelhamid, T. Alioto, I. Antoshechkin, M. T. Baer, P. Batut, I. Bell, K. Bell, S. Chakrabortty, X. Chen, J. Chrast, J. Curado, T. Derrien, J. Drenkow, E. Dumais, J. Dumais, R. Duttagupta, M. Fastuca, K. Fejes-Toth, P. Ferreira, S. Foissac, M. J. Fullwood, H. Gao, D. Gonzalez, A. Gordon, H. P. Gunawardena, C. Howald, S. Jha, R. Johnson, P. Kapranov, B. King, C. Kingswood, G. Li, O. J. Luo, E. Park, J. B. Preall, K. Presaud, P. Ribeca, B. A. Risk, D. Robyr, X. Ruan, M. Sammeth, K. S. Sandhu, L. Schaeffer, L.-H. See, A. Shahab, J. Skancke, A. M. Suzuki, H. Takahashi, H. Tilgner, D. Trout, N. Walters, H. Wang, J. Wrobel, Y. Yu, Y. Hayashizaki, J. Harrow, M. Gerstein, T. J. Hubbard, A. Reymond, S. E. Antonarakis, G. J. Hannon, M. C. Giddings, Y. Ruan, B. Wold, P. Carninci, R. Guigó, T. R. Gingeras, K. R. Rosenbloom, C. A. Sloan, K. Learned, V. S. Malladi, M. C. Wong, G. P. Barber, M. S. Cline, T. R. Dreszer, S. G. Heitner, D. Karolchik, W. J. Kent, V. M. Kirkup, L. R. Meyer, J. C. Long, M. Maddren, B. J. Raney, T. S. Furey, L. Song, L. L. Grasfeder, P. G. Giresi, B.-K. Lee, A. Battenhouse, N. C. Sheffield, J. M. Simon, K. A. Showers, A. Safi, D. London, A. A. Bhinge, C. Shestak, M. R. Schaner, S. Ki Kim, Z. Z. Zhang, P. A. Mieczkowski, J. O. Mieczkowska, Z. Liu, R. M. McDaniell, Y. Ni, N. U. Rashid, M. J. Kim, S. Adar, Z. Zhang, T. Wang, D. Winter, D. Keefe, E. Birney, V. R. Iyer, J. D. Lieb, G. E. Crawford, G. Li, K. S. Sandhu, M. Zheng, P. Wang, O. J. Luo, A. Shahab, M. J. Fullwood, X. Ruan, Y. Ruan, R. M. Myers, F. Pauli, B. A. Williams, J. Gertz, G. K. Marinov, T. E. Reddy, J. Vielmetter, E. Partridge, D. Trout, K. E. Varley, C. Gasper, The ENCODE Project Consortium, Overall coordination (data analysis coordination), Data production leads (data production), Lead analysts (data analysis), Writing group, NHGRI project management (scientific management), Principal investigators (steering committee), Boise State University and University of North Carolina at Chapel Hill Proteomics groups (data production and analysis), Broad Institute Group (data production and analysis), U. of G. Cold Spring Harbor Center for Genomic Regulation, Barcelona, RIKEN, Sanger Institute, University of Lausanne, Genome Institute of Singapore group (data production and analysis), Data coordination center at UC Santa Cruz (production data coordination), E. Duke University University of Texas, Austin, University of North Carolina-Chapel Hill group (data production and analysis), Genome Institute of Singapore group (data production and analysis), C. HudsonAlpha Institute UC Irvine, Stanford group (data production and analysis), An integrated encyclopedia of DNA elements in the human genome. Nature 489, 57–74 (2012).

9. J. S. Mattick, I. V. Makunin, Non-coding RNA. Hum. Mol. Genet. 15, R17–R29 (2006).

10. T. R. Cech, J. A. Steitz, The Noncoding RNA Revolution—Trashing Old Rules to Forge New Ones. Cell 157, 77–94 (2014).

11. A. Isakova, D. D. Liu, I. Cvijovic, R. Sinha, A. E. Eastman, S. Saul, A. Detweiler, N. Neff, S. Einav, I. L. Weissman, S. R. Quake, Beyond polyA: scalable single-cell total RNA-seq unifies coding and non-coding transcriptomics. bioRxiv [Preprint] (2025). 10.1101/2025.08.08.669394.

12. D. W. McKellar, M. Mantri, M. M. Hinchman, J. S. L. Parker, P. Sethupathy, B. D. Cosgrove, I. De Vlaminck, Spatial mapping of the total transcriptome by in situ polyadenylation. Nat. Biotechnol. 41, 513–520 (2023).

13. G. Crowley, S. R. Quake, Benchmarking cell type and gene set annotation by large language models with AnnDictionary. Nat. Commun. 16, 9511 (2025).

14. T. Derrien, R. Johnson, G. Bussotti, A. Tanzer, S. Djebali, H. Tilgner, G. Guernec, D. Martin, A. Merkel, D. G. Knowles, J. Lagarde, L. Veeravalli, X. Ruan, Y. Ruan, T. Lassmann, P. Carninci, J. B. Brown, L. Lipovich, J. M. Gonzalez, M. Thomas, C. A. Davis, R. Shiekhattar, T. R. Gingeras, T. J. Hubbard, C. Notredame, J. Harrow, R. Guigó, The GENCODE v7 catalog of human long noncoding RNAs: Analysis of their gene structure, evolution, and expression. Genome Res. 22, 1775–1789 (2012).

15. C.-C. Hon, J. A. Ramilowski, J. Harshbarger, N. Bertin, O. J. L. Rackham, J. Gough, E. Denisenko, S. Schmeier, T. M. Poulsen, J. Severin, M. Lizio, H. Kawaji, T. Kasukawa, M. Itoh, A. M. Burroughs, S. Noma, S. Djebali, T. Alam, Y. A. Medvedeva, A. C. Testa, L. Lipovich, C.-W. Yip, I. Abugessaisa, M. Mendez, A. Hasegawa, D. Tang, T. Lassmann, P. Heutink, M. Babina, C. A. Wells, S. Kojima, Y. Nakamura, H. Suzuki, C. O. Daub, M. J. L. de Hoon, E. Arner, Y. Hayashizaki, P. Carninci, A. R. R. Forrest, An atlas of human long non-coding RNAs with accurate 5′ ends. Nature 543, 199–204 (2017).

16. M. N. Cabili, C. Trapnell, L. Goff, M. Koziol, B. Tazon-Vega, A. Regev, J. L. Rinn, Integrative annotation of human large intergenic noncoding RNAs reveals global properties and specific subclasses. Genes Dev. 25, 1915–1927 (2011).

17. S. J. Liu, M. A. Horlbeck, S. W. Cho, H. S. Birk, M. Malatesta, D. He, F. J. Attenello, J. E. Villalta, M. Y. Cho, Y. Chen, M. A. Mandegar, M. P. Olvera, L. A. Gilbert, B. R. Conklin, H. Y. Chang, J. S. Weissman, D. A. Lim, CRISPRi-based genome-scale identification of functional long noncoding RNA loci in human cells. Science 355, eaah7111 (2017).

18. K. A. Dittmar, J. M. Goodenbour, T. Pan, Tissue-specific differences in human transfer RNA expression. PLoS Genet. 2, e221 (2006).

19. O. Pinkard, S. McFarland, T. Sweet, J. Coller, Quantitative tRNA-sequencing uncovers metazoan tissue-specific tRNA regulation. Nat. Commun. 11, 4104 (2020).

20. J. Cavaillé, K. Buiting, M. Kiefmann, M. Lalande, C. I. Brannan, B. Horsthemke, J.-P. Bachellerie, J. Brosius, A. Hüttenhofer, Identification of brain-specific and imprinted small nucleolar RNA genes exhibiting an unusual genomic organization. Proc. Natl. Acad. Sci. 97, 14311–14316 (2000).

21. É. Fafard-Couture, D. Bergeron, S. Couture, S. Abou-Elela, M. S. Scott, Annotation of snoRNA abundance across human tissues reveals complex snoRNA-host gene relationships. Genome Biol. 22, 172 (2021).

22. A. Isakova, T. Fehlmann, A. Keller, S. R. Quake, A mouse tissue atlas of small noncoding RNA. Proc. Natl. Acad. Sci. 117, 25634–25645 (2020).

23. J. A. Oo, K. Pálfi, T. Warwick, I. Wittig, C. Prieto-Garcia, V. Matkovic, I. Tomašković, F. Boos, J. Izquierdo Ponce, T. Teichmann, K. Petriukov, S. Haydar, L. Maegdefessel, Z. Wu, M. D. Pham, J. Krishnan, A. H. Baker, S. Günther, H. D. Ulrich, I. Dikic, M. S. Leisegang, R. P. Brandes, Long non-coding RNA *PCAT19* safeguards DNA in quiescent endothelial cells by preventing uncontrolled phosphorylation of RPA2. Cell Rep. 41, 111670 (2022).

24. K. R. Mitchelson, W.-Y. Qin, Roles of the canonical myomiRs miR-1, –133 and – 206 in cell development and disease. World J. Biol. Chem. 6, 162–208 (2015).

25. E. Goñi, A. M. Mas, J. Gonzalez, A. Abad, M. Santisteban, P. Fortes, M. Huarte, M. Hernaez, Uncovering functional lncRNAs by scRNA-seq with ELATUS. Nat. Commun. 15, 9709 (2024).

26. I. Yanai, H. Benjamin, M. Shmoish, V. Chalifa-Caspi, M. Shklar, R. Ophir, A. Bar-Even, S. Horn-Saban, M. Safran, E. Domany, D. Lancet, O. Shmueli, Genome-wide midrange transcription profiles reveal expression level relationships in human tissue specification. Bioinforma. Oxf. Engl. 21, 650–659 (2005).

27. H. Li, J. Janssens, M. D. Waegeneer, S. S. Kolluru, K. Davie, V. Gardeux, W. Saelens, F. P. A. David, M. Brbić, K. Spanier, J. Leskovec, C. N. McLaughlin, Q. Xie, R. C. Jones, K. Brueckner, J. Shim, S. G. Tattikota, F. Schnorrer, K. Rust, T. G. Nystul, Z. Carvalho-Santos, C. Ribeiro, S. Pal, S. Mahadevaraju, T. M. Przytycka, A. M. Allen, S. F. Goodwin, C. W. Berry, M. T. Fuller, H. White-Cooper, E. L. Matunis, S. DiNardo, A. Galenza, L. E. O’Brien, J. A. T. Dow, F. C. A. Consortium§, H. Jasper, B. Oliver, N. Perrimon, B. Deplancke, S. R. Quake, L. Luo, S. Aerts, D. Agarwal, Y. Ahmed-Braimah, M. Arbeitman, M. M. Ariss, J. Augsburger, K. Ayush, C. C. Baker, T. Banisch, K. Birker, R. Bodmer, B. Bolival, S. E. Brantley, J. A. Brill, N. C. Brown, N. A. Buehner, X. T. Cai, R. Cardoso-Figueiredo, F. Casares, A. Chang, T. R. Clandinin, S. Crasta, C. Desplan, A. M. Detweiler, D. B. Dhakan, E. Donà, S. Engert, S. Floc’hlay, N. George, A. J. González-Segarra, A. K. Groves, S. Gumbin, Y. Guo, D. E. Harris, Y. Heifetz, S. L. Holtz, F. Horns, B. Hudry, R.-J. Hung, Y. N. Jan, J. S. Jaszczak, G. S. X. E. Jefferis, J. Karkanias, T. L. Karr, N. S. Katheder, J. Kezos, A. A. Kim, S. K. Kim, L. Kockel, N. Konstantinides, T. B. Kornberg, H. M. Krause, A. T. Labott, M. Laturney, R. Lehmann, S. Leinwand, J. Li, J. S. S. Li, K. Li, K. Li, L. Li, T. Li, M. Litovchenko, H.-H. Liu, Y. Liu, T.-C. Lu, J. Manning, A. Mase, M. Matera-Vatnick, N. R. Matias, C. E. McDonough-Goldstein, A. McGeever, A. D. McLachlan, P. Moreno-Roman, N. Neff, M. Neville, S. Ngo, T. Nielsen, C. E. O’Brien, D. Osumi-Sutherland, M. N. Özel, I. Papatheodorou, M. Petkovic, C. Pilgrim, A. O. Pisco, C. Reisenman, E. N. Sanders, G. dos Santos, K. Scott, A. Sherlekar, P. Shiu, D. Sims, R. V. Sit, M. Slaidina, H. E. Smith, G. Sterne, Y.-H. Su, D. Sutton, M. Tamayo, M. Tan, I. Tastekin, C. Treiber, D. Vacek, G. Vogler, S. Waddell, W. Wang, R. I. Wilson, M. F. Wolfner, Y.-C. E. Wong, A. Xie, J. Xu, S. Yamamoto, J. Yan, Z. Yao, K. Yoda, R. Zhu, R. P. Zinzen, Fly Cell Atlas: A single-nucleus transcriptomic atlas of the adult fruit fly. Science, doi: 10.1126/science.abk2432 (2022).

28. Z. Yao, C. T. J. van Velthoven, M. Kunst, M. Zhang, D. McMillen, C. Lee, W. Jung, J. Goldy, A. Abdelhak, M. Aitken, K. Baker, P. Baker, E. Barkan, D. Bertagnolli, A. Bhandiwad, C. Bielstein, P. Bishwakarma, J. Campos, D. Carey, T. Casper, A. B. Chakka, R. Chakrabarty, S. Chavan, M. Chen, M. Clark, J. Close, K. Crichton, S. Daniel, P. DiValentin, T. Dolbeare, L. Ellingwood, E. Fiabane, T. Fliss, J. Gee, J. Gerstenberger, A. Glandon, J. Gloe, J. Gould, J. Gray, N. Guilford, J. Guzman, D. Hirschstein, W. Ho, M. Hooper, M. Huang, M. Hupp, K. Jin, M. Kroll, K. Lathia, A. Leon, S. Li, B. Long, Z. Madigan, J. Malloy, J. Malone, Z. Maltzer, N. Martin, R. McCue, R. McGinty, N. Mei, J. Melchor, E. Meyerdierks, T. Mollenkopf, S. Moonsman, T. N. Nguyen, S. Otto, T. Pham, C. Rimorin, A. Ruiz, R. Sanchez, L. Sawyer, N. Shapovalova, N. Shepard, C. Slaughterbeck, J. Sulc, M. Tieu, A. Torkelson, H. Tung, N. Valera Cuevas, S. Vance, K. Wadhwani, K. Ward, B. Levi, C. Farrell, R. Young, B. Staats, M.-Q. M. Wang, C. L. Thompson, S. Mufti, C. M. Pagan, L. Kruse, N. Dee, S. M. Sunkin, L. Esposito, M. J. Hawrylycz, J. Waters, L. Ng, K. Smith, B. Tasic, X. Zhuang, H. Zeng, A high-resolution transcriptomic and spatial atlas of cell types in the whole mouse brain. Nature 624, 317–332 (2023).

29. D. O’Reilly, M. Dienstbier, S. A. Cowley, P. Vazquez, M. Drozdz, S. Taylor, W. S. James, S. Murphy, Differentially expressed, variant U1 snRNAs regulate gene expression in human cells. Genome Res. 23, 281–291 (2013).

30. P. Vazquez-Arango, D. O’Reilly, Variant snRNPs: New players within the spliceosome system. RNA Biol. 15, 17–25 (2017).

31. J. W. Mabin, P. W. Lewis, D. A. Brow, H. Dvinge, Human spliceosomal snRNA sequence variants generate variant spliceosomes. RNA 27, 1186–1203 (2021).

32. P. Landgraf, M. Rusu, R. Sheridan, A. Sewer, N. Iovino, A. Aravin, S. Pfeffer, A. Rice, A. O. Kamphorst, M. Landthaler, C. Lin, N. D. Socci, L. Hermida, V. Fulci, S. Chiaretti, R. Foà, J. Schliwka, U. Fuchs, A. Novosel, R.-U. Müller, B. Schermer, U. Bissels, J. Inman, Q. Phan, M. Chien, D. B. Weir, R. Choksi, G. D. Vita, D. Frezzetti, H.-I. Trompeter, V. Hornung, G. Teng, G. Hartmann, M. Palkovits, R. D. Lauro, P. Wernet, G. Macino, C. E. Rogler, J. W. Nagle, J. Ju, F. N. Papavasiliou, T. Benzing, P. Lichter, W. Tam, M. J. Brownstein, A. Bosio, A. Borkhardt, J. J. Russo, C. Sander, M. Zavolan, T. Tuschl, A Mammalian microRNA Expression Atlas Based on Small RNA Library Sequencing. Cell 129, 1401–1414 (2007).

33. D. P. Bartel, MicroRNAs: Target Recognition and Regulatory Functions. Cell 136, 215–233 (2009).

34. D. P. Bartel, Metazoan MicroRNAs. Cell 173, 20–51 (2018).

35. W.-W. Liang, S. Müller, S. Hart, H.-H. Wessels, A. Méndez-Mancilla, A. Sookdeo, O. Choi, C. Caragine, A. Corman, L. Lu, O. Kolumba, B. Williams, N. Sanjana, Transcriptome-scale RNA-targeting CRISPR screens reveal essential lncRNAs in human cells. figshare [Preprint] (2025). 10.6084/m9.figshare.30370171.v1.

36. A. R. Buxbaum, G. Haimovich, R. H. Singer, In the right place at the right time: visualizing and understanding mRNA localization. Nat. Rev. Mol. Cell Biol. 16, 95–109 (2015).

37. S. Das, M. Vera, V. Gandin, R. H. Singer, E. Tutucci, Intracellular mRNA transport and localized translation. Nat. Rev. Mol. Cell Biol. 22, 483–504 (2021).

38. E. Lécuyer, H. Yoshida, N. Parthasarathy, C. Alm, T. Babak, T. Cerovina, T. R. Hughes, P. Tomancak, H. M. Krause, Global analysis of mRNA localization reveals a prominent role in organizing cellular architecture and function. Cell 131, 174–187 (2007).

39. D. E. Henze, A. P. Tsai, T. Wyss-Coray, S. R. Quake, Simultaneous spatial transcriptomics and morphology profiling as tools to explore how microglia change with age. *Nat*. Aging, 1–17 (2026).

40. E. T. Wang, J. M. Taliaferro, J.-A. Lee, I. P. Sudhakaran, W. Rossoll, C. Gross, K. R. Moss, G. J. Bassell, Dysregulation of mRNA Localization and Translation in Genetic Disease. J. Neurosci. 36, 11418–11426 (2016).

41. O. J. Ziff, J. Harley, Y. Wang, J. Neeves, G. Tyzack, F. Ibrahim, M. Skehel, A. M. Chakrabarti, G. Kelly, R. Patani, Nucleocytoplasmic mRNA redistribution accompanies RNA binding protein mislocalization in ALS motor neurons and is restored by VCP ATPase inhibition. Neuron 111, 3011–3027.e7 (2023).

42. M. S. Fernandopulle, J. Lippincott-Schwartz, M. E. Ward, RNA transport and local translation in neurodevelopmental and neurodegenerative disease. Nat. Neurosci. 24, 622–632 (2021).

43. M. C. Bridges, A. C. Daulagala, A. Kourtidis, LNCcation: lncRNA localization and function. J. Cell Biol. 220, e202009045 (2021).

44. M. Trabucchi, R. Mategot, Subcellular Heterogeneity of the microRNA Machinery. Trends Genet. TIG 35, 15–28 (2019).

45. F. M. Fazal, S. Han, K. R. Parker, P. Kaewsapsak, J. Xu, A. N. Boettiger, H. Y. Chang, A. Y. Ting, Atlas of Subcellular RNA Localization Revealed by APEX-Seq. Cell 178, 473–490.e26 (2019).

46. F. Zhou, P. Tan, S. Liu, L. Chang, J. Yang, M. Sun, Y. Guo, Y. Si, D. Wang, J. Yu, Y. Ma, Subcellular RNA distribution and its change during human embryonic stem cell differentiation. Stem Cell Rep. 19, 126–140 (2024).

47. J. Carlevaro-Fita, R. Johnson, Global Positioning System: Understanding Long Noncoding RNAs through Subcellular Localization. Mol. Cell 73, 869–883 (2019).

48. X. Dai, Y. Li, W. Liu, X. Pan, C. Guo, X. Zhao, J. Lv, H. Lei, L. Zhang, Application of RNA subcellular fraction estimation method to explore RNA localization regulation. G3 GenesGenomesGenetics 12, jkab371 (2021).

49. K. Chatterjee, R. T. Nostramo, Y. Wan, A. K. Hopper, tRNA dynamics between the nucleus, cytoplasm and mitochondrial surface: Location, location, location. Biochim. Biophys. Acta 1861, 373–386 (2018).

50. A. K. Hopper, R. T. Nostramo, tRNA Processing and Subcellular Trafficking Proteins Multitask in Pathways for Other RNAs. Front. Genet. 10 (2019).

51. U. Fischer, C. Englbrecht, A. Chari, Biogenesis of spliceosomal small nuclear ribonucleoproteins. WIREs RNA 2, 718–731 (2011).

52. J. Park, S. H. Ahn, M. G. Shin, H. K. Kim, S. Chang, tRNA-Derived Small RNAs: Novel Epigenetic Regulators. Cancers 12 (2020).

53. T. Hu, C. Lu, Y. Xia, L. Wu, J. Song, C. Chen, Q. Wang, Small nucleolar RNA SNORA71A promotes epithelial-mesenchymal transition by maintaining ROCK2 mRNA stability in breast cancer. Mol. Oncol. 16, 1947–1965 (2022).

54. H.-W. Hwang, E. A. Wentzel, J. T. Mendell, A Hexanucleotide Element Directs MicroRNA Nuclear Import. Science 315, 97–100 (2007).

55. A. F. Palazzo, E. S. Lee, Non-coding RNA: what is functional and what is junk? Front. Genet. 6 (2015).

56. M. Kapur, M. J. Molumby, C. Guzman, S. Heinz, S. L. Ackerman, Cell-type-specific expression of tRNAs in the brain regulates cellular homeostasis. Neuron 112, 1397–1415.e6 (2024).

57. L. Gao, A. Behrens, G. Rodschinka, S. Forcelloni, S. Wani, K. Strasser, D. D. Nedialkova, Selective gene expression maintains human tRNA anticodon pools during differentiation. Nat. Cell Biol. 26, 100–112 (2024).

58. M. V. Rodnina, W. Wintermeyer, Fidelity of aminoacyl-tRNA selection on the ribosome: kinetic and structural mechanisms. Annu. Rev. Biochem. 70, 415–435 (2001).

59. M. A. Sørensen, C. G. Kurland, S. Pedersen, Codon usage determines translation rate in *Escherichia coli*. J. Mol. Biol. 207, 365–377 (1989).

60. A. Dana, T. Tuller, The effect of tRNA levels on decoding times of mRNA codons. Nucleic Acids Res. 42, 9171–9181 (2014).

61. I. Frumkin, M. J. Lajoie, C. J. Gregg, G. Hornung, G. M. Church, Y. Pilpel, Codon usage of highly expressed genes affects proteome-wide translation efficiency. Proc. Natl. Acad. Sci. 115, E4940–E4949 (2018).

62. S. Rudorf, Efficiency of protein synthesis inhibition depends on tRNA and codon compositions. PLOS Comput. Biol. 15, e1006979 (2019).

63. W. Qian, J.-R. Yang, N. M. Pearson, C. Maclean, J. Zhang, Balanced Codon Usage Optimizes Eukaryotic Translational Efficiency. PLOS Genet. 8, e1002603 (2012).

64. W. Gao, C. J. Gallardo-Dodd, C. Kutter, Cell type–specific analysis by single-cell profiling identifies a stable mammalian tRNA–mRNA interface and increased translation efficiency in neurons. Genome Res. 32, 97–110 (2022).

65. M. S. Kowalczyk, I. Tirosh, D. Heckl, T. N. Rao, A. Dixit, B. J. Haas, R. K. Schneider, A. J. Wagers, B. L. Ebert, A. Regev, Single-cell RNA-seq reveals changes in cell cycle and differentiation programs upon aging of hematopoietic stem cells. Genome Res. 25, 1860–1872 (2015).

66. A. I. Tirosh, B. Izar, S. M. Prakadan, M. H. Wadsworth, D. Treacy, J. J. Trombetta, A. Rotem, C. Rodman, C. Lian, G. Murphy, M. Fallahi-Sichani, K. Dutton-Regester, J.-R. Lin, O. Cohen, P. Shah, D. Lu, A. S. Genshaft, T. K. Hughes, C. G. K. Ziegler, S. W. Kazer, A. Gaillard, K. E. Kolb, A.-C. Villani, C. M. Johannessen, A. Y. Andreev, E. M. Van Allen, M. Bertagnolli, P. K. Sorger, R. J. Sullivan, K. T. Flaherty, D. T. Frederick, J. Jané-Valbuena, C. H. Yoon, O. Rozenblatt-Rosen, A. K. Shalek, A. Regev, L. A. Garraway, Dissecting the multicellular ecosystem of metastatic melanoma by single-cell RNA-seq. Science 352, 189–196 (2016).

66. E. Z. Macosko, A. Basu, R. Satija, J. Nemesh, K. Shekhar, M. Goldman, I. Tirosh, A. R. Bialas, N. Kamitaki, E. M. Martersteck, J. J. Trombetta, D. A. Weitz, J. R. Sanes, A. K. Shalek, A. Regev, S. A. McCarroll, Highly Parallel Genome-wide Expression Profiling of Individual Cells Using Nanoliter Droplets. Cell 161, 1202–1214 (2015).

67. A. Hernandez-Segura, T. V. de Jong, S. Melov, V. Guryev, J. Campisi, M. Demaria, Unmasking Transcriptional Heterogeneity in Senescent Cells. Curr. Biol. CB 27, 2652–2660.e4 (2017).

68. D. Saul, R. L. Kosinsky, E. J. Atkinson, M. L. Doolittle, X. Zhang, N. K. LeBrasseur, R. J. Pignolo, P. D. Robbins, L. J. Niedernhofer, Y. Ikeno, D. Jurk, J. F. Passos, L. J. Hickson, A. Xue, D. G. Monroe, T. Tchkonia, J. L. Kirkland, J. N. Farr, S. Khosla, A new gene set identifies senescent cells and predicts senescence-associated pathways across tissues. Nat. Commun. 13, 4827 (2022).

70. A. G. S. Kinker, A. C. Greenwald, R. Tal, Z. Orlova, M. S. Cuoco, J. M. McFarland, A. Warren, C. Rodman, J. A. Roth, S. A. Bender, B. Kumar, J. W. Rocco, P. A. C. M. Fernandes, C. C. Mader, H. Keren-Shaul, A. Plotnikov, H. Barr, A. Tsherniak, O. Rozenblatt-Rosen, V. Krizhanovsky, S. V. Puram, A. Regev, I. Tirosh, Pan-cancer single-cell RNA-seq identifies recurring programs of cellular heterogeneity. Nat. Genet. 52, 1208–1218 (2020).

69. M. Kitagawa, K. Kitagawa, Y. Kotake, H. Niida, T. Ohhata, Cell cycle regulation by long non-coding RNAs. Cell. Mol. Life Sci. CMLS 70, 4785–4794 (2013).

70. K. Abdelmohsen, M. Gorospe, Noncoding RNA Control of Cellular Senescence. Wiley Interdiscip. Rev. RNA 6, 615–629 (2015).

71. N. Romero-Barrios, M. F. Legascue, M. Benhamed, F. Ariel, M. Crespi, Splicing regulation by long noncoding RNAs. Nucleic Acids Res. 46, 2169–2184 (2018).

72. G. Guiducci, L. Stojic, Long Noncoding RNAs at the Crossroads of Cell Cycle and Genome Integrity. Trends Genet. 37, 528–546 (2021).

73. S. Ghafouri-Fard, T. Khoshbakht, B. M. Hussen, A. Baniahmad, W. Branicki, M. Taheri, A. Eghbali, Emerging Role of Non-Coding RNAs in Senescence. Front. Cell Dev. Biol. 10, 869011 (2022).

74. T. M. Geiman, K. Muegge, Lsh, an SNF2/helicase family member, is required for proliferation of mature T lymphocytes. Proc. Natl. Acad. Sci. 97, 4772–4777 (2000).

75. R. Mjelle, S. A. Hegre, P. A. Aas, G. Slupphaug, F. Drabløs, P. Saetrom, H. E. Krokan, Cell cycle regulation of human DNA repair and chromatin remodeling genes. DNA Repair 30, 53–67 (2015).

76. F. Lessard, S. Igelmann, C. Trahan, G. Huot, E. Saint-Germain, L. Mignacca, N. Del Toro, S. Lopes-Paciencia, B. Le Calvé, M. Montero, X. Deschênes-Simard, M. Bury, O. Moiseeva, M.-C. Rowell, C. E. Zorca, D. Zenklusen, L. Brakier-Gingras, V. Bourdeau, M. Oeffinger, G. Ferbeyre, Senescence-associated ribosome biogenesis defects contributes to cell cycle arrest through the Rb pathway. Nat. Cell Biol. 20, 789–799 (2018).

77. Y. Cheng, S. Wang, H. Zhang, J.-S. Lee, C. Ni, J. Guo, E. Chen, S. Wang, A. Acharya, T.-C. Chang, M. Buszczak, H. Zhu, J. T. Mendell, A non-canonical role for a small nucleolar RNA in ribosome biogenesis and senescence. Cell 187, 4770–4789.e23 (2024).

78. M. Gorospe, K. Abdelmohsen, MicroRegulators come of age in senescence. Trends Genet. TIG 27, 233–241 (2011).

79. M. Deschênes, B. Chabot, The emerging role of alternative splicing in senescence and aging. Aging Cell 16, 918–933 (2017).

80. Z. Ding, G. Ma, B. Zhou, S. Cheng, W. Tang, Y. Han, L. Chen, W. Pang, Y. Chen, D. Yang, H. Cao, Targeting miR-29 mitigates skeletal senescence and bolsters therapeutic potential of mesenchymal stromal cells. Cell Rep. Med. 5 (2024).

81. K. Abdelmohsen, A. Panda, M.-J. Kang, J. Xu, R. Selimyan, J.-H. Yoon, J. L. Martindale, S. De, W. H. Wood, K. G. Becker, M. Gorospe, Senescence-associated lncRNAs: senescence-associated long noncoding RNAs. Aging Cell 12, 890–900 (2013).

82. P. K. Puvvula, LncRNAs Regulatory Networks in Cellular Senescence. Int. J. Mol. Sci. 20, 2615 (2019).

83. M. Martin, Cutadapt removes adapter sequences from high-throughput sequencing reads. EMBnet.journal 17, 10–12 (2011).

84. B. Langmead, S. L. Salzberg, Fast gapped-read alignment with Bowtie 2. Nat. Methods 9, 357–359 (2012).

85. J. M. Mudge, S. Carbonell-Sala, M. Diekhans, J. G. Martinez, T. Hunt, I. Jungreis, J. E. Loveland, C. Arnan, I. Barnes, R. Bennett, A. Berry, A. Bignell, D. Cerdán-Vélez, K. Cochran, L. T. Cortés, C. Davidson, S. Donaldson, C. Dursun, R. Fatima, M. Hardy, P. Hebbar, Z. Hollis, B. T. James, Y. Jiang, R. Johnson, G. Kaur, M. Kay, R. J. Mangan, M. Maquedano, L. M. Gómez, N. Mathlouthi, R. Merritt, P. Ni, E. Palumbo, T. Perteghella, F. Pozo, S. Raj, C. Sisu, E. Steed, D. Sumathipala, M.-M. Suner, B. Uszczynska-Ratajczak, E. Wass, Y. T. Yang, D. Zhang, R. D. Finn, M. Gerstein, R. Guigó, T. J. P. Hubbard, M. Kellis, A. Kundaje, B. Paten, M. L. Tress, E. Birney, F. J. Martin, A. Frankish, GENCODE 2025: reference gene annotation for human and mouse. Nucleic Acids Res. 53, D966–D975 (2025).

86. P. P. Chan, T. M. Lowe, GtRNAdb 2.0: an expanded database of transfer RNA genes identified in complete and draft genomes. Nucleic Acids Res. 44, D184–D189 (2016).

87. A. Kozomara, M. Birgaoanu, S. Griffiths-Jones, miRBase: from microRNA sequences to function. Nucleic Acids Res. 47, D155–D162 (2019).

88. B. Kaminow, D. Yunusov, A. Dobin, STARsolo: accurate, fast and versatile mapping/quantification of single-cell and single-nucleus RNA-seq data. bioRxiv [Preprint] (2021). 10.1101/2021.05.05.442755.

89. F. A. Wolf, P. Angerer, F. J. Theis, SCANPY: large-scale single-cell gene expression data analysis. Genome Biol. 19, 15 (2018).

90. S. J. Fleming, M. D. Chaffin, A. Arduini, A.-D. Akkad, E. Banks, J. C. Marioni, A. A. Philippakis, P. T. Ellinor, M. Babadi, Unsupervised removal of systematic background noise from droplet-based single-cell experiments using CellBender. Nat. Methods 20, 1323–1335 (2023).

91. S. L. Wolock, R. Lopez, A. M. Klein, Scrublet: Computational Identification of Cell Doublets in Single-Cell Transcriptomic Data. Cell Syst. 8, 281–291.e9 (2019).

92. I. Korsunsky, N. Millard, J. Fan, K. Slowikowski, F. Zhang, K. Wei, Y. Baglaenko, M. Brenner, P. Loh, S. Raychaudhuri, Fast, sensitive and accurate integration of single-cell data with Harmony. Nat. Methods 16, 1289–1296 (2019).

93. A. Isakova, N. Neff, S. R. Quake, Single-cell quantification of a broad RNA spectrum reveals unique noncoding patterns associated with cell types and states. Proc. Natl. Acad. Sci. 118, e2113568118 (2021).

